# SPITFIR(e): A supermaneuverable algorithm for restoring 2D-3D fluorescence images and videos, and background subtraction

**DOI:** 10.1101/2022.01.04.474883

**Authors:** Sylvain Prigent, Hoai-Nam Nguyen, Ludovic Leconte, Cesar Augusto Valades-Cruz, Bassam Hajj, Jean Salamero, Charles Kervrann

## Abstract

While fluorescent microscopy imaging has become the spearhead of modern biology as it is able to generate long-term videos depicting 4D nanoscale cell behaviors, it is still limited by the optical aberrations and the photon budget available in the specimen and to some extend to photo-toxicity. A direct consequence is the necessity to develop flexible and “off-road” algorithms in order to recover structural details and improve spatial resolution, which is critical when pushing the illumination to the low levels in order to limit photo-damages. Moreover, as the processing of very large temporal series of images considerably slows down the analysis, special attention must be paid to the feasibility and scalability of the developed restoration algorithms. To address these specifications, we present a very flexible method designed to restore 2D-3D+Time fluorescent images and subtract undesirable out-of-focus background. We assume that the images are sparse and piece-wise smooth, and are corrupted by mixed Poisson-Gaussian noise. To recover the unknown image, we consider a novel convex and non-quadratic regularizer Sparse Hessian Variation) defined as the mixed norms which gathers image intensity and spatial second-order derivatives. This resulting restoration algorithm named SPITFIR(e) (SParse fIT for Fluorescence Image Restoration) utilizes the primal-dual optimization principle for energy minimization and can be used to process large images acquired with varied fluorescence microscopy modalities. It is nearly parameter-free as the practitioner needs only to specify the amount of desired sparsity (weak, moderate, high). Experimental results in lattice light sheet, stimulated emission depletion, multifocus microscopy, spinning disk confocal, and wide-field microscopy demonstrate the generic ability of the SPITFIR(e) algorithm to efficiently reduce noise and blur, and to subtract undesirable fluorescent background, while avoiding the emergence of deconvolution artifacts.

---

Fluorescence microscopy provides a very powerful framework to biologists for observing, analyzing, and studying specific fluorescently-tagged structures and biological processes at very high spatial and temporal resolutions. In general, the specimen is first labelled by fluorescent molecules before being excited by the illumination light of a given wavelength; upon the excitation, fluorophore re-emits light of relatively longer wavelength which is then collected by photosensitive sensors to form the digitized image of the observed sample. Despite number of advantages, there are two major limitations of fluorescence microscopy. The first limitation is the presence of photon (shot) noise in acquired images. Shot noise is mainly due to the quantum nature of light, implying that the arrival of a photon on a sensor is a random event and thus the number of incident photons over a period of time is a random variable depending on the brightness of the light source. Moreover, experimentally in biological samples the signal-to-noise-ratio (SNR) is usually very low because low dose of illumination light is required to avoid photo-bleaching (i.e., progressive fading of the emission intensity) of fluorescent molecules and to preserve specimen integrity (photo-toxicity) ^1;2^. Additionally, the quality of acquired fluorescence images is worsened by the blurring effects induced by several factors such as excitation wavelength, immersion medium refraction, specimen thickness, and the limited aperture of the objective which results in light diffraction through the optical system. The diffraction phenomenon implies that light emitted by an infinitely small point source appears wider at the focal plan and spreads into a specific pattern called “point spread function” (PSF).

Noise and blur not only degrade the image quality in terms of overall visualization but also have a negative influence on specimen analysis, including detection and segmentation of biological structures. To improve the quality of images acquired by fluorescence microscopes, restoration (deconvolution and/or denoising) is then frequently applied as pre-processing step before quantitative analysis, and amounts to recovering the original unknown image *u* from the observed noisy and blurred image *f* represented as follows: *f* = 𝒯 (*h* * *u*), where *h* denotes the 2D or 3D spatial response of the device representing the blur related to the optical system (e.g., PSF) assumed to be linear shift-invariant, *\** denotes the convolution operator, and 𝒯 is a degradation operator modeling the measurement noise. If the image is not blurred but mainly corrupted by noise, *h* is represented by the Dirac (or impulse) function and the restoration problem translates into a denoising problem. Finally, the degradation operator (shot noise and readout noise) is usually modeled as a mixed Poisson-Gaussian distribution which is non-stationary and signal-dependent.

In the last twenty years, many restoration methods have been investigated in order to “inverse” the model in 2D-3D fluorescence microscopy. The most popular restoration approaches are linear methods (e.g., Wiener filtering). Despite the simplicity and low-computation-requirement, they usually do not restore fine image details at frequency components that are beyond the bandwidth of the PSF (i.e., the support of its Fourier transform). Another popular technique is the Richardson-Lucy (RL) algorithm ^3;4;5^. While well-dedicated to Poisson noise removal, RL deconvolution creates structural noise and artifacts after a small number of iterations, which constitutes a major problem in fluorescence imaging^6^. Actually, the most flexible and robust deconvolution methods consist in minimizing an energy functional composed of a data fidelity term and a regularization term that encompasses prior knowledge (positivity, smoothness, sparsity,…) about the solution. The seminal Tikhonov-Miller (TM) approach ^7;8^ may be considered as the starting point to the development of regularization methods in fluorescence microscopy. While it was successful in many applications, the TM regularizer tends to produce blurred images since low gradient magnitudes are encouraged in the entire image, including at image discontinuities. To avoid over-smoothing caused by quadratic functionals, non-quadratic regularizers have been first studied, especially the Total Variation (TV) regularizer that penalizes the *L*_1_ norm of the gradient. Nevertheless, TV creates “stair-casing” effects, which are particular undesirable in fluorescence imaging^10;11^. To overcome this disadvantage, Lefkimmiatis *et al*. ^12^ recently proposed a family of convex regularizers built upon the matrix norm of the Hessian (Schatten norm), computed at each point of the image. This piece-wise smoothness promoting regularizer combined with a non-quadratic data term, was recently evaluated in fluorescence microscopy^13^. Nevertheless, this regularizer does not impose sparsity prior on the fluorescent signals which is also an important feature in fluorescence imaging. This idea was investigated by Arigovindan *et al*. ^14^ who reported good results 3D wide-field fluorescence images by using a non-convex sparsity-promoting regularizer; this regularizer (named here Log Sparse Hessian Variation (LHSV)) has been implemented into the ER-Decon deconvolution algorithm. Concurrently, other sparsity-promoting methods, focused on first-order differentiation-based regularizers have been investigated in medical imaging such as GraphNet (GN) ^15;16;17^ and Sparse Variation (SV) ^18^, and in image processing (e.g., TV-*L*_1_ ^19;20^). Nevertheless, the sparsity-promoting regularizers reported in Table 1 have limitations. They were designed for 2D medical imaging^15;16;17;18^ or 3D microscopy^14;21^ but never for both, are based on the first-order ^15;16;17;18^ or second derivatives ^14;21^, are formulated as non-convex^14^ or convex ^21^ optimization problems which are now efficiently solved by performant algorithms such as ADMM^22;23^ and FISTA^24^.

**Table 1:**
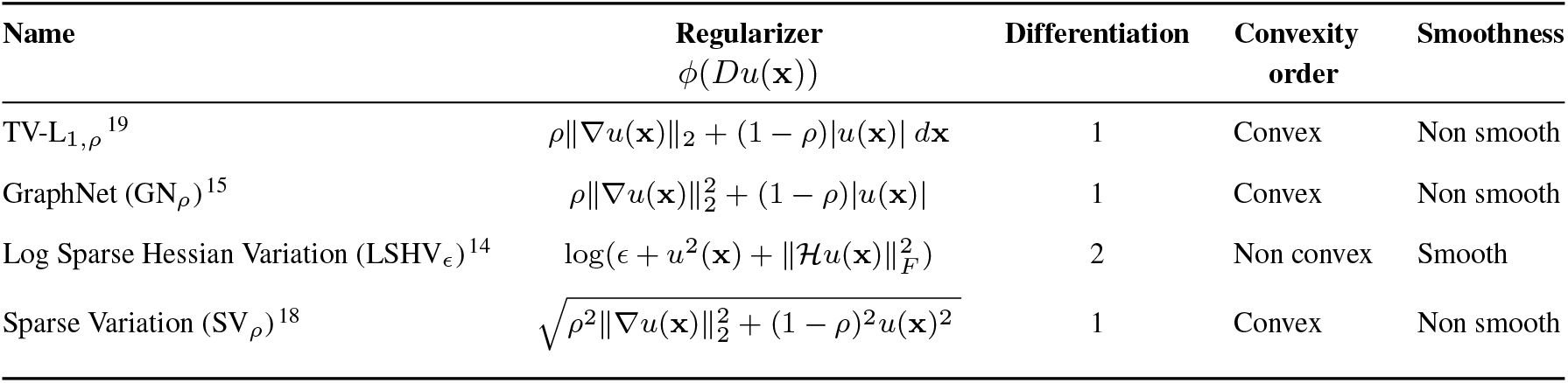
Properties of regularizers used for restoring fluorescence microscopy images.

**Table 2:**
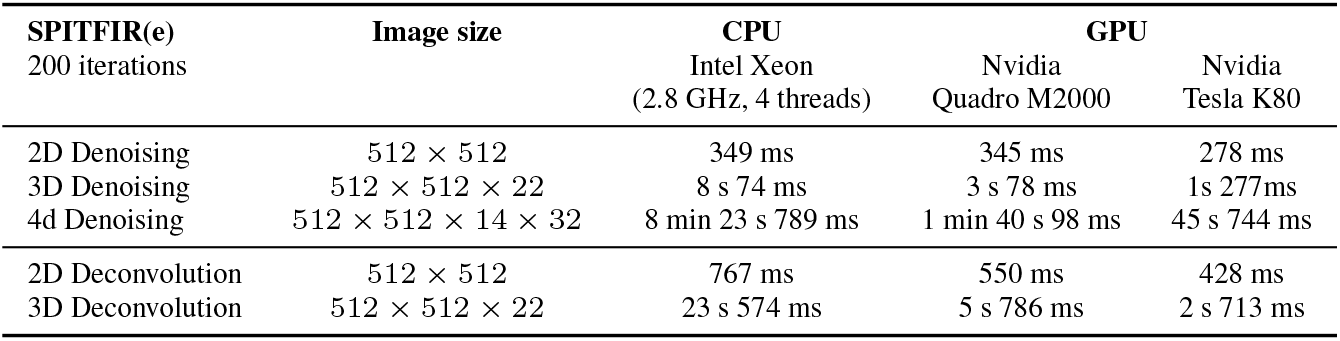
Computing times of SPITFIR(e) (200 iterations for energy minimization).

To overcome the main limitations of previous methods, we introduce a flexible method (SPITFIR(e)) to deconvolve and/or denoise a large range of 2D/3D fluorescence microscopy images and videos (Supplementary Fig. 1). SPITFIR(e) is able to handle low signal-to-noise ratios, high-dimensional images (2D/3D/4D multispectral images), and characteristics of microscopy set-ups, especially if the PSF is not well known. The resulting approach includes a novel 2D/3D/4D sparsity promoting regularizer, can adapt to most if not all fluorescence techniques without fine-tuning of parameters, and shows robustness to different sources of noise and degradations. The SPITFIR(e) parameters are automatically adjusted from the image contents and the practitioner needs only to specify if the target image is “highly sparse”, “moderately sparse” or “weakly sparse”. We validated the performance of our approach and compared SPITFIR(e) to competitive deconvolution methods on several datasets, and we confirmed the results on real images acquired with lattice light sheet (LLSM), stimulated emission depletion (STED), multi-focus (MFM), spinning disk confocal (CM), and wide-field microscopy (WF). For the evaluation on different imaging modes, we have chosen to apply SPITFIR(e) on a limited number of structures relatively difficult to resolve or for which the photon budget is critical, i.e., mitochondria, microtubules and biomolecules in motion in living cells.

## Results

### Overview of SPITFIR(e)

We present SPITFIR(e), an algorithm well grounded in the regularization theory for inverse problems, that can robustly recover information from general multidimensional noisy and blurry fluorescence images. This algorithm is able to adapt to several if not all microscopy modalities as illustrated in Figs. 1-3, and to different sources of noise and blur. The practitioner needs only to set the desired level of sparsity, which amounts here to selecting if the image is *“highly sparse”, “moderately sparse”* or *“weakly sparse”*. A “weakly” sparse image contains clutter and complex contents. Unlike the traditional methods requiring the tedious manual adjustment of the parameter that balances the data fidelity term and the prior term, SPITFIR(e) includes a self-tuning and scale-invariant technique for automatic adaptation (Methods). The minimization of the sum of the two convex terms is originally implemented in a computationally efficient way with a recent version of proximal-splitting optimization algorithm ^25;26^ (Methods and Supplementary Notes (Appendix B)). In what follows, we show that SPITFIR(e) can restore 3D fluorescence images under low excitation light excitation conditions by chaining denoising and deconvolution with the same algorithm with very small computing times compared to other competitive methods ^13;21^. A “plane-by-plane” (PbyP) deconvolution strategy is also included for fast deconvolution if the 3D PSF is not well known. It turns out that the decomposition into a series of 2D decoupled deconvolution problems provided very good results in a small computing time. The nearly parameter-free SPITFIR(e) algorithm has been implemented both in CPU and GPU, is interfaced with Napari (https://www.napari-hub.org/plugins/napari-sdeconv) and can be downloaded for free here: https://github.com/sylvainprigent/simglib.

**Fig. 1:**
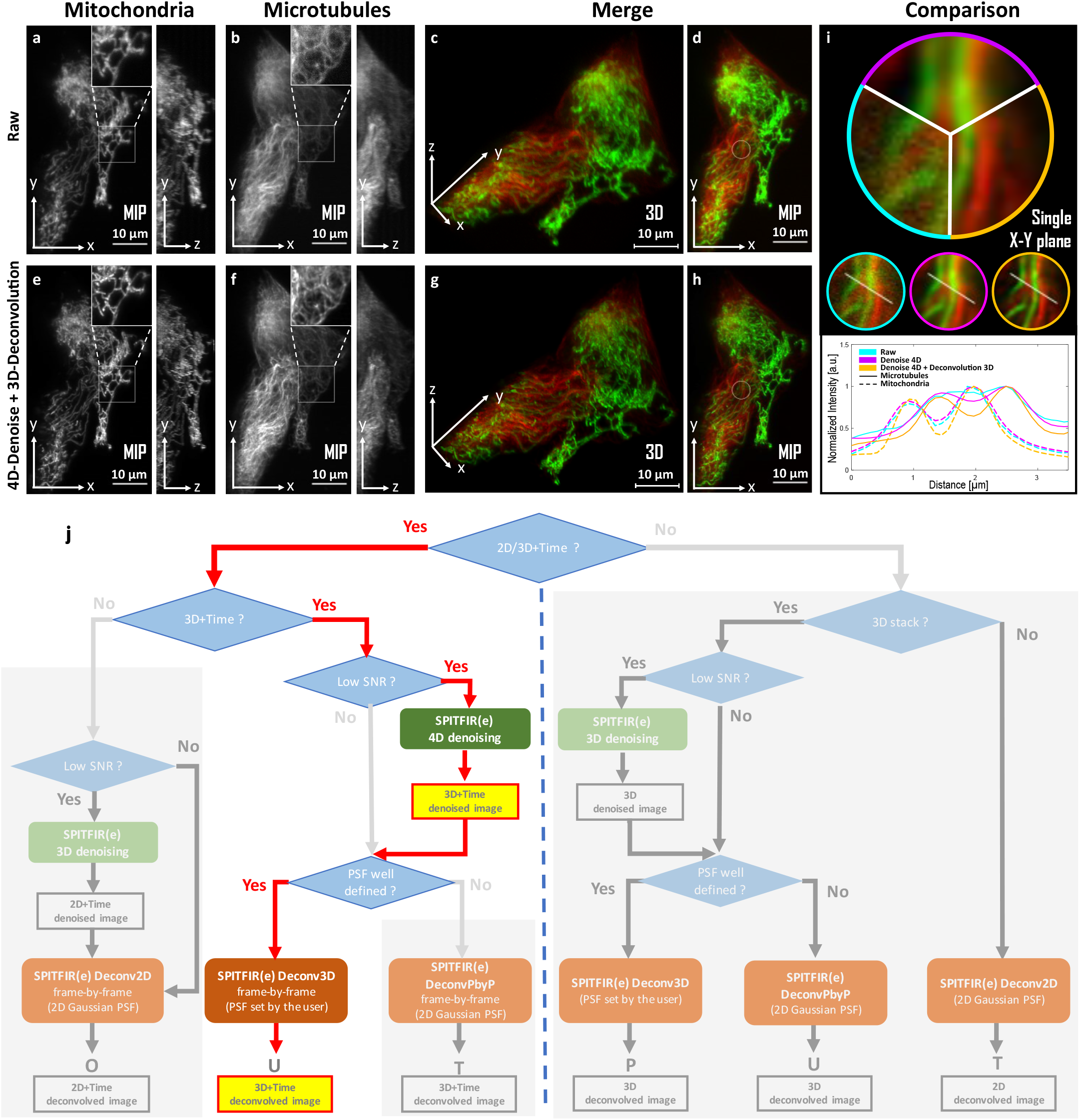
SPITFIR(e) for 4D-denoising, background subtraction, and 3D-deconvolution on LLSM images. 60 planes 3D volumes of live RPE1 cells double stained with PKMR for Mitochondria (**a**,**e**) and with Tubulin Tracker^*TM*^ Deep Red for Microtubules (**b**,**f**) were acquired within 2.2*s* per stack using lattice light-sheet microscopy. MIP of representative raw (see Material and Methods section) images for Mitochondria and Microtubules, respectively, before (**a**,**b**) and after (**e, f**) SPITFIR(e) 4D denoising and 3D deconvolution (3D Gaussian PSF, *σ*_*xy*_ = 1.5 pixels and *σ*_*z*_ = 1.0 pixel). Insets are zoomed area illustrating SPITFIR(e) improvement in spatial resolution and SNR. 3D angular views and MIP of composite images before (**c**,**d**) and after (**g**,**h**) SPITFIR(e) treatment (Red for Microtubules; Green for Mitochondria). 4D denoising SPITFIR(e) image improvement shown on a single x-y plane (**i**) zoomed from the inset indicated in (**d, h**). Raw image is indicated as a blue lined sector in the upper part and in the left circle, beneath; 4D denoised image is similarly indicated in magenta, while the 4D denoised + 3D deconvolved image is in yellow. Intensity line profiles were measured as indicated in the 3 circles and plotted for each processing step and both Mitochondria and Microtubules at the lower part of (**i**). **j**, Workflow for image restoration extracted from the global flow chart in Supplementary Fig. 1. Scale bars are indicated in the bottom right corner (**a-h**). Related videos are shown as Supplementary movies S1 and S2.

**Fig. 2:**
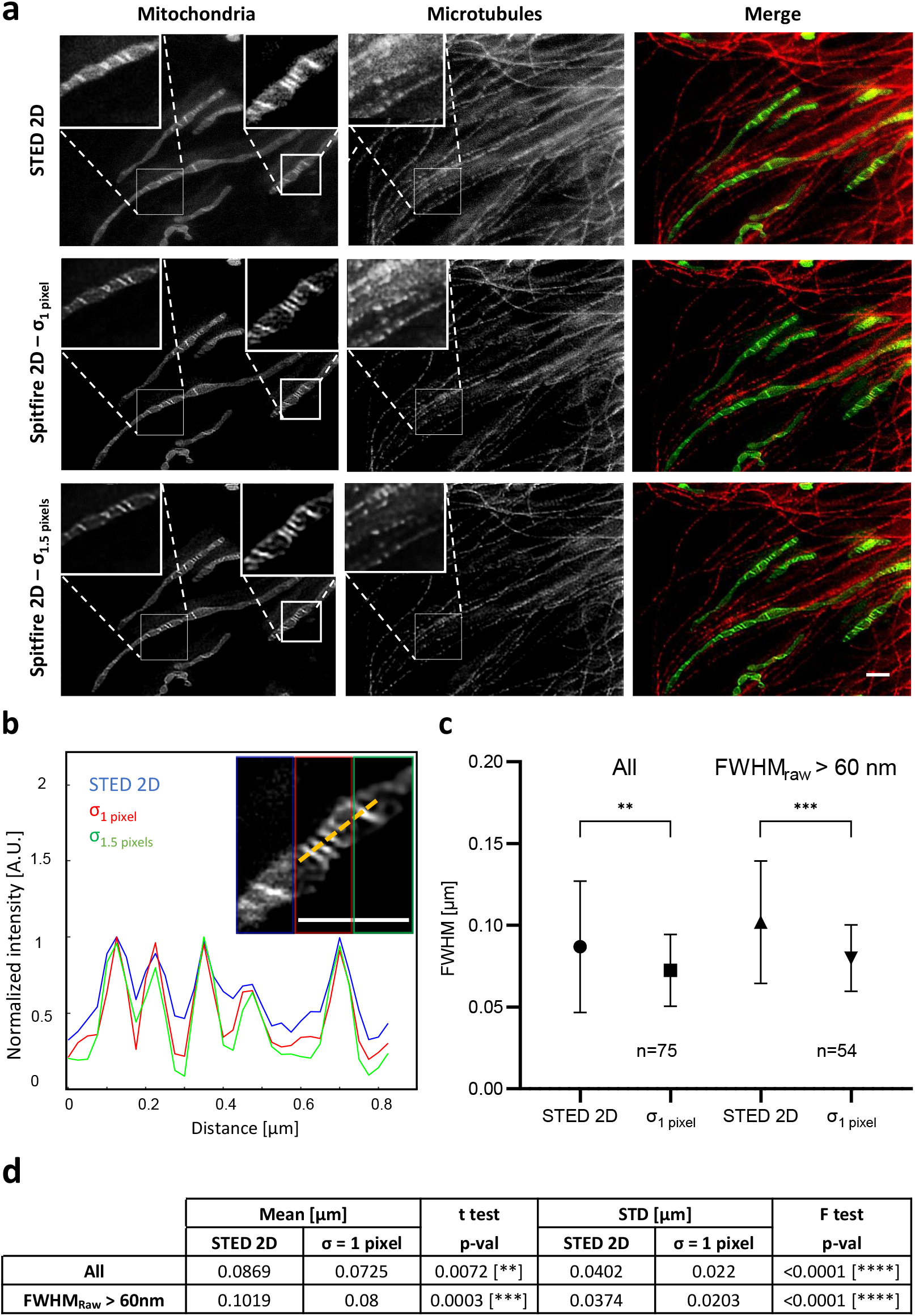
SPITFIR(e) increases the signal-to-noise of low-light images in STED imaging. Live RPE1 cells double stained with PKMO (mitochondria) and Tubulin TrackerTM Deep Red (microtubules) were imaged by STED nanoscopy before and after being processed by SPITFIR(e) (**a**). Partial overview of 2D STED raw data of RPE1 cells for mitochondria, microtubules and their superimposed images are shown from left to right (**a**, upper panel). 2D STED images were improved by SPITFIR(e) using two different 2D Gaussian model *σ*_*xy*_ = 1.0 pixel (**a**, middle panel) and *σ*_*xy*_ = 1.5 pixels (**a**, lower panel). Insets (1 and 2) show zoomed area where SPITFIR(e) image quality improvement is illustrated by comparison of two processing conditions (middle and lower panel) with the same area in the raw STED image. Inset(**b**) shows a magnified composite image where the 2D STED raw part is indicated as a blue lined rectangle and 2D denoised + 2D deconvolved parts are indicated in red (*σ*_*xy*_ = 1.0 pixel) and green (*σ*_*xy*_ = 1.5 pixels). (**b**) Normalized intensity line profiles were measured in the insets (**b**) for cristae regions. Yellow line indicated in the inset (**b5**) serve to identify fluorescence profiles. Pixel size is equal to 25 nm. Scale bars: 1 *μ*m in (**a**) and (**b**). Line profiles of individual cristae (n=75, from 8 different acquisitions) were fitted using a Gaussian model. Full width half maximum (FWHM) was estimated on raw STED and 2D denoised + 2D deconvolved with *σ*_*xy*_ = 1.0 pixel (**c**). SPITFIR(e) improvement is shown separately for all the line profiles analyzed and the line profiles from bigger cristae (FWHM_*raw*_ = 60 nm), indication of lower SNR. Statistics analysis is including in (**d**). It indicates the improvement of average resolution (unpaired two-sample *t*-test) and variance of cristae resolution (*F* -test).

**Fig. 3:**
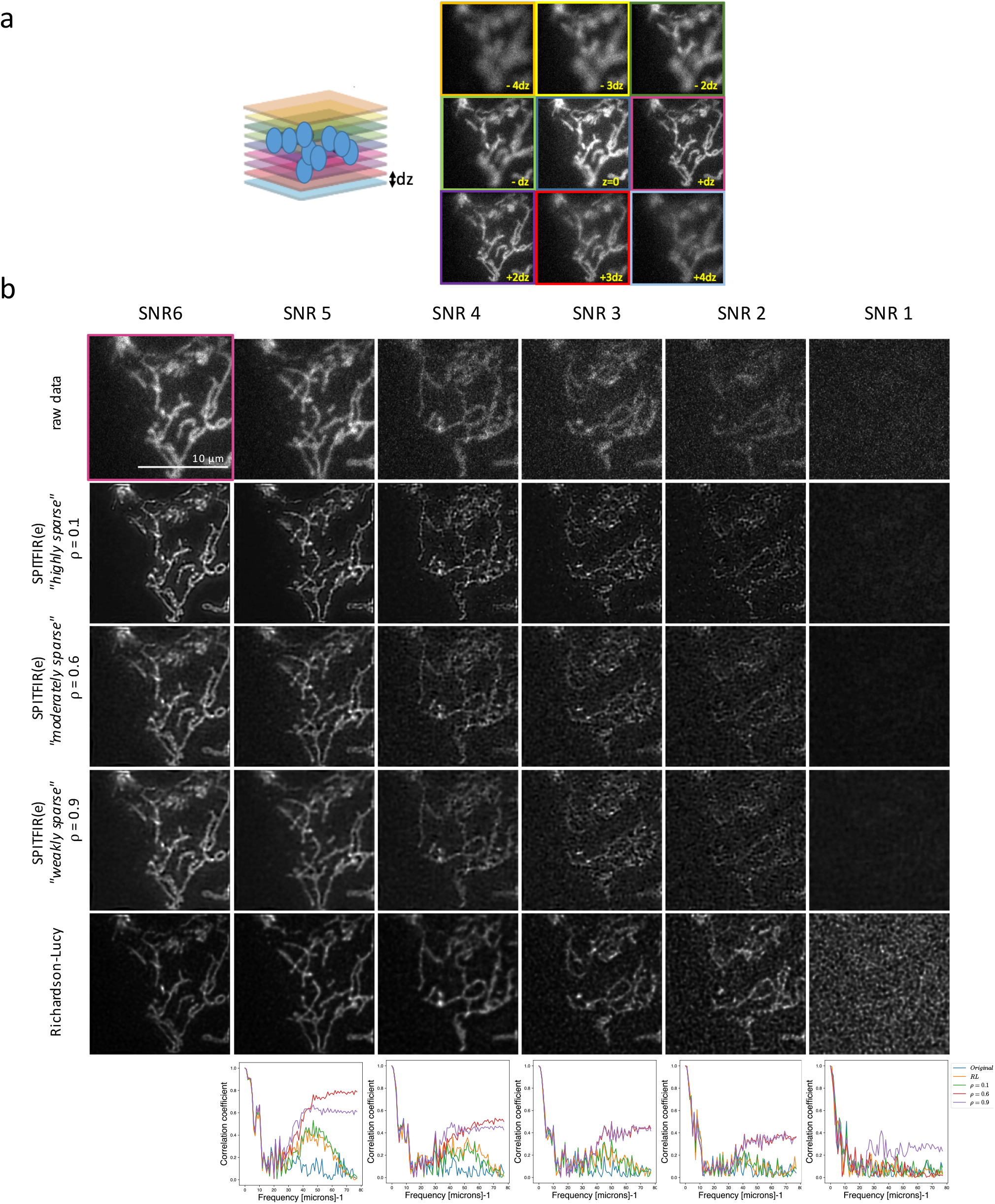
A comparison of wide-field multifocus microscopy image reconstruction with different amounts of sparsity and SNR values. (**a**) MFM allows simultaneous acquisition of a 3D stack of nine images equally spaced with (dz = 330 nm) focal 2D images (dx, dy = 120 nm) with a frame rate up to 100 images per second, thus covering the whole volume of mitochondria in a single 3D image ^31^. The set-up is based on a custom diffraction grating that forms multiple focus-shifted images followed by a chromatic correction module. (**b**) The image grid display the deconvolution results on the 6th plane for each SNR value (+dz in **a**)) of the wide-field MFM stack depicting mitochondria in U2OS cells transfected with TOM20 (translocase of outer mitochondrial membrane) fused to GFP (GFP-TOM20). The 3D stack has been deconvolved with 3D Gaussian PSF model (*σ*_*xy*_ = 1.5 pixels and *σ*_*z*_ = 1.5 pixel) and with three different levels of sparsity (“weak”, “moderate”, and “high”). The plots in the bottom are the Fourier Shell Correlation of the reconstructed image for each SNR value, by considering the higher SNR value (SNR 6) as reference image. Scale bar: 10 *μ*m.

### SPITFIR(e) is flexible and adapts to every fluorescence microscopy modality

SPITFIR(e) includes several strategies to adapt to microscopy specificities, and to particular spatial and temporal acquisition conditions (Supplementary Fig. 1). For instance, the practitioner can apply the conventional 3D deconvolution strategy if the PSF is perfectly known. If this is not the case, the PbyP strategy which consists in separately deconvolving each XY section of the 3D stack given a 2D Gaussian PSF model, can be applied. Surprisingly, the PbyP strategy could produce better visual results and is faster than direct 3D deconvolution with an imperfect or approximately measured 3D PSF. Meanwhile, in the case of low-photon regimes or low-exposure times, the results may be significantly improved by applying the “denoise-before-deconvolve” approach. For instance, a 3D+Time (4D) volume can be denoised as a whole, while the planes of each 3D stack are independently deconvolved with a 2D Gaussian PSF model. An exemplary demonstration of the flexibility of SPITFIR(e) is given by the experiments performed with a lattice light sheet microscope ^27^ (LLSM) which allows to acquire 3D images for several hours with limited photobleaching. In LLSM imaging, a sheet of light inclined about 32 degrees illuminate one slice of the sample while an objective oriented at 90 degrees from the illumination slice allows to observe and image the illuminated slice. Moving the illumination slice allows then to create a 3D image. As the illumination angle makes the raw image stack skewed, post-processing is required to create a 3D “deskewed” stack. Consequently, deconvolving 3D LLSM images is not straightforward as the PSF is not well characterized and the deskew operation affects the noise statistics. We then design the following SPITFIR(e) deconvolution pipeline according to the flowchart illustrated in Supplementary Fig. 1. We hypothesized here that chaining the denoising and deconvolution steps could provide better restorations results. So, first we denoise the 3D+Time image sequence (4D denoising) with SPITFIR(e) (the PSF function *h* is a Dirac function). Then we deconvolve each 3D stack separately with SPITFIR(e) by using an anisotropic 3D Gaussian PSF model. The settings of the PSF are chosen given the image resolution along the XY and Z axes.

We illustrate the efficiency of the “denoise-before-deconvolve” approach using mitochondria (Fig. 1). Live imaging mitochondria and their interactions with other intracellular components at high spatial resolution in 3D and high temporal frequency, is known to be challenging. Beyond the usual optical limits, labeled mitochondria are very sensitive to light ^28^, which could induce artifactual fluorescence signals such as exaltation, probes dissociation and finally fragmentation of mitochondria and cell death. This may impair accurate 3D+Time observation of their behaviors in normal and stress conditions. For this reasons, imaging mitochondria dynamics and structural features will serve as one of the biological thread in the following parts of the manuscript. We here linked 3D LLSM at low illumination regimes to efficiently preserve fluorescence from photobleaching and consequently from phototoxic effects, with 4D denoising and 3D image deconvolution (3D Gaussian PSF, *σ*_*xy*_ = 1.5 pixels and *σ*_*z*_ = 1.0 pixel) by using SPITFIR(e). We imaged live RPE1 cells in full volume (Fig. 1 labeled for mitochondria (Fig. 1**(a)** and **(e)**) and microtubules (Fig. 1**(b)** and **(f)**). A reconstituted comparison of the successive steps of the SPITFIR(e) workflow is shown in Fig. 1**(i)**. For the sake of visual clarity, only one single plane of the zoomed area of a stack is shown, (as indicated in Fig. 1**(d)** and **(h))** and clock wise decomposed from the raw image to full processing. More visual representations can be appreciated in supplementary Movies 1 and 2, in 3D+Time along both XY and Z axes. Interestingly, chaining denoising and deconvolution with SPITFIR(e) is particularly visible on the microtubule staining, while improvement in resolution is more obvious on mitochondria (compare insets of zoomed area between Fig. 1**(b)** and **(f)** and between (**a**) and **(e)**). Line intensity plots through the center of the fluorescent filamentous structure demonstrate the recovering of the signal from the noise (Fig. 1**(i)** bottom part). Furthermore, the potential of SPITFIR(e) in LLSM imaging is also illustrated on a volume of 58 planes depicting endosomes stained with the AP2 proteins (Supplementary Fig. 2) known to be involved in the early steps of endocytosis ^29^. In this experiment, we compared both the 3D deconvolution and PbyP deconvolution strategies with a 3D Gaussian model (*σ*_*xy*_ = 1.0 pixel and *σ*_*z*_ = 0.8 pixel) and a 2D Gaussian model (*σ*_*xy*_ = 1.0 pixel), respectively. The line intensity plots (**b**) through the center of the fluorescent spots demonstrates that SPITFIR(e) with the BbyP strategy preserves the intensity peaks, and better narrows the full-width-half-maximum (FWMH) when compared to RL^3^ and HR-C3D^13^. These two case-studies indicate the ability of SPITFIR(e) to recover general shapes (i.e., filaments and point-like objects) in noisy and blurred fluorescence images.

Meanwhile, we also evaluated the flexibility of SPITFIR(e) on spinning disk confocal microscopy images. Spinning disk microscopes with many pinholes allows a high speed acquisition of images and therefore is compatible with acquisition regimes corresponding to live cell imaging. Nevertheless, time exposure must be reduced to capture fast intracellular events, such as biomolecules dynamics. Supplementary Fig. 3 shows the maximum intensity projection along the Z-axis of a stack depicting Rab5 proteins stained with Green Fluorescence Protein (GFP) close to endosomes within a cell. The image is blurred and noisy, and contains small spots at the resolution limit of the microscope against a dark background. Supplementary Fig. 4 shows the maximum intensity projection along the Z-axis of a stack depicting mCherry-actin filaments, that is thin tubular-like structures. This image is apparently not as sparse as the other image. These two examples are challenging because the PSF is approximately known and the image contents are very different. The PSF is assumed to be approximated by a 3D and a 2D Gaussian models. In each figure, we reported the results obtained with four strategies (3D deconvolution, PbyP deconvolution, 3D denoising + 3D deconvolution, 3D denoising + PbyP deconvolution) and different levels of sparsity. Applying denoising beforehand enables here to better enhance weak fluorescent signals while removing blur and background. From Supplementary Figs. 3-4, it turns out that the images are visually best results on the GFP-Rab5 and mCherry-actin images when the sparsity is “high” and “moderate”, respectively

With SPITFIR(e), we have also improved the resolution limits of current STED microscopy. This nanoscopy is one of the very few light microscopy techniques allowing to solve mitochondria cristae organization, as illustrated here in live cells (Fig. 2). STED microscopy provides subdiffraction resolution while preserving useful aspects of fluorescence microscopy, such as optical sectioning, and molecular specificity and sensitivity. The PSF shape depends on several factors such as optics, laser power, fluorescence response of the specimen, and thermal drift which can add deformation by sheering. In Fig. 2, we deconvolved with SPITFIR(e) two 2D images depicting mitochondria and their alignment on microtubules, by applying SPITFIR(e) 2D with a 2D Gaussian PSF model and two different bandwidths *σ*_*xy*_ of size 1.5 pixels and 1.0 pixel respectively, respectively (Fig. 2**(a)**). We selected two regions of interest (ROI) in the mitochondria channel to better appreciate the restoration results in (Fig. 2(**a, b**)). Normalized fluorescence intensity line profiles, positioned at two distinct cristae regions of mitochondria (Fig. 2(**b**)) show the dual improvement of SPITFIR(e) with both an increase in the SNR and to a lower extend, an enhanced lateral resolution. Two simulated PSFs sizes were proposed to evaluate the capacity of SPITFIR(e) to improve the image resolution. The first PSF (*σ*_*xy*_ = 1.0 pixel) was similar than the expected experimental PSF of our microscope and the second PSF (*σ*_*xy*_ = 1.5 pixels) corresponds to the situation where there is an issue of defocusing. Clearly in both cases we observe an image improvement of the two individual cristae line profiles. The fitting of a Gaussian model to intensity profiles (*n* = 75, from 8 different acquisitions) allowed us to estimate the resolution at FWHM of raw and restored images (**b-c**). The obtained resolution is statistically improved when SPITFIR(e) is applied (with a smaller variance), as reported in **d**, and agrees well with the reported electron microscopy results ^30^.

### SPITFIR(e) reliably restores weak fluorescent signals under extreme low-light conditions

Additional demonstrations of the ability of SPITFIR(e) to retrieve the high spatial-frequency information under extreme light conditions in 3D multifocus microscopy and spinning disk confocal microscopy, are presented below. Here, we focus on the performance limits of 3D deconvolution knowing that it is possible to improve the results by denoising the images beforehand as described above.

First, we tested SPITFIR(e) to visualize the 3D dynamics of mitochondria in living cells under extreme low-light conditions with multifocus (MFM) for different SNR values by reducing the excitation light dose (see Fig. 3(**a**)). In this experiment, each 3D stack was restored with a 3D Gaussian PSF model. The results on 2D plane (6th plane in (**a**)) are shown in Fig. 3 for six different SNRs values using the same sample. We considered three different levels of sparsity: “highly sparse”, “moderately sparse”, and “weakly sparse”. In Fig. 3(**b**) (bottom), we reported the corresponding Fourier Shell Correlation curves computed by selecting the image with the higher SNR value (SNR 6) as reference image. We can notice that for SNR 5 we get a good correlation in high frequency and the restored images are visually very similar to images with SNR 6. For SNR 4 to SNR 2, we can observe that SPITFIR(e) gradually looses correlation in high frequencies, and the reconstructed images are visually less accurate. For very poor SNR conditions (SNR 1), the images are very noisy and SPITFIR(e) (3D deconvolution) was not able to recover the signals and the high frequencies correlation is close to zero. Meanwhile, the images obtained by applying RL deconvolution did not allow us to retrieve information at SNR 4 and below. Moreover, artifacts and spatial distortions appear in the final image reconstructed when using RL deconvolution. Considering a sparse-promoting regularization helped here reveal structural details at lower SNR values. Overall, the most satisfying results with SPITFIR(e) are obtained with “moderate” sparsity.

Next, we evaluated the performance of SPITFIR(e) on 3D spinning disk confocal images acquired with different speeds and excitation light doses to capture mast movements and reduce photobleaching and photo-toxicity, respectively. To assess the overall sensitivity of SPITFIR(e), 4D images were captured at variable excitation light levels, reducing light intensity by a factor of 2 and 4. When we reduce by half the laser power (raw data denoted *I*_1*/*2_ in Supplementary Fig. 5), the bleaching effect is less visible and the SNR is lower. Furthermore, when we reduce the laser power by factor 4 (raw data denoted *I*_1*/*4_ in Supplementary Fig. 5) we do not observe bleaching effect anymore. In Supplementary Fig. 5 (**b**), the three time-lapse temporal series display reduced bleaching curves which correlates with the excitation light intensity levels and SNR values. Nevertheless, the reduction factor has the undesirable consequence of lowering the signal-to-noise ratio (SNR) of the image, resulting in a poor-quality image which is also degraded by optics. We applied the proposed SPITFIR(e) restoration confocal pipeline on the three 3D+Time images *I, I*_1*/*2_, *I*_1*/*4_. The results shown in Fig. 5 are named SPITFIR(e)(*I*), SPITFIR(e)(*I*_1*/*2_), SPITFIR(e)(*I*_1*/*4_). We can notice that even with a low SNR (*I*_1*/*4_) we obtain a good enough quality restored images for spot detection. We conclude that that even at low illumination doses and low SNR, we can obtain a good enough quality restored images with SPITFIR(e), which is helpful to preserve the integrity of samples, and till get exploitable data for quantitative analysis.

### SPITFIR(e) quantitatively outperforms state-of-the-art algorithms

In fluorescence microscopy, restoration algorithms were dedicated to remove Poisson-Gaussian noise originating from the low-photons regimes (Poisson noise) and dark current induced by the electronic imaging detectors (Gaussian noise). In Methods (and Supplementary Notes (Appendix A)), we formally demonstrate that a quadratic fidelity term combined with any regularization term is optimal to handle Poisson-Gaussian noise. This new result allows us to apply SPITFIR(e) on images acquired with low light doses (Fig. 3 and Supplementary Fig. 5) and to quantitatively compare state-of-the-art deconvolution algorithms mainly developed for Gaussian noise removal, and applied here to fluorescence microscopy images.

First, we fairly benchmarked the performance of a large collection of competitive deconvolution algorithms given in Table 1, including the popular deconvolution methods such as RL, iterative constrained Tikhonov-Miller (ICTM), Gold-Meinel (GM)) ^32^, and applied on artificially noisy and blurred 2D images. We considered four ground-truth images shown in Supplementary Table 2 which were blurred with 2D Gaussian PSF model with different standard deviation values *σ*_*xy*_. A Gaussian noise with zero mean and variance *τ*^2^ was also added to these images in order to generate observed noisy and blurry data. In Supplementary Table 2, we reported the best possible PSNR values for each method (with optimal parameter adjustment) and the true PSF size *σ*_*xy*_ and noise variance *τ*^2^. In this experiment, RL and GM algorithm were often ranked at the end of the benchmark, while SPITFIR(e) provided the best PSNR values in most of cases. In terms of visual assessment, we noticed noise amplification with RL (Fig. 6(**c**) and GM (Fig. 6 (**d**), resulting in the apparition of undesired high-intensity pixels (i.e., the “night sky” effect) and unrealistic reconstructed structures since the high-frequency components were not correctly restored. ICTM produced deconvolution results without noise amplification artifacts but tends to over-smooth image details due to the quadratic nature of Tikhonov-Miller penalty, while TV (**f**), TV-*L*_1_ (**g**) and SV (**l**) regularizers generate artificial sharper edges (stair-casing effects), respectively. HV (**m**) generates visually more pleasant deconvolution results. Regarding smooth-approximation based regularizations, LSHV (**j**) provide results with restored details which are slightly sharper than those obtained with ICTM (**e**), but noise is not sufficiently removed. Finally, SPITFIR(e) (SHV, **n**) avoids over-smoothing and over-sharpening effect while preserving details and removing noise in the background. These quantitative results were confirmed on 3D microscopy images by considering additional performance criteria.

As the implementation in 3D of most of aforementioned regularizers in Table 1 is not always possible, we focused on the Hessian-regularized deconvolution algorithm ^13^ (HR-C3D) as it was demonstrated to outperform previous software such as Huygens, Microvolution ^33^, and ER-Decon (LSHV) ^14^. RL deconvolution is also considered and serves here as a baseline method. A first comparison of SPITFIR(e), HR-C3D and RL applied to LLSM imaging (see Supplementary Fig. 2) has been previously described. For the sake of objectivity, we present here the results obtained on a three-channel volume (22 planes) depicting actin (green channel, stained with Alexa Fluor 488 phalloidin), nucleus (blue challenge, stained with DAPI) and mitochondria (red channel, stained with MitoTracker Red CMXRos), acquired with a wide-field microscope and already used for algorithm evaluation by Ikoma *et al*. ^13^. In Supplementary Fig. 7, we report the normalized root-mean-square error (NRMSE), global Pearson correlation coefficients (RSP), PSNR values, and Fourier Shell Correlation (FSC) curves ^14^, between the restored wide-field images obtained with the two competing methods (SPITFIR(e), HR-C3D) and the reference spinning disk confocal image deconvolved with the Huygens software ^13^, for each channel. We carefully followed the recommendations given in ref ^13^ to fairly normalize the deconvolved images with respect to the *L*_2_ norm of the deconvolved confocal image (with offset correction). The good performance of SPITFIR(e) is first confirmed by the PSNR/NMSE values which are slightly higher/smaller than those obtained with HR-C3D ^13^. The resolution estimated from FSC curves is also higher (nucleus, actin) or comparable (mitochondria). This is especially demonstrative at high frequencies in the case of actin filaments and cell nucleus. SPITFIR(e) produces visually smoother images than HR-C3D but more correlated to the spinning disk confocal microscopy images

Finally, we conducted experiments on a 3D confocal microscopy and compared the results of RL, SPITFIR(e) and HR-C3D to those produced by CARE ^34^, a supervised deep-learning-based deconvolution method (Supplementary Fig. 8). As CARE is learned from a training set of dedicated biological structures, we evaluate the algorithms on the stack provided by the authors ^34^ and depicting the envelopes of nuclei stained with GFP-LAP2b (developing eye of zebrafish (Daniorerio) embryos. We focused on the nuclear envelop as this channel contains more structural details. In Supplementary Figs. 8 we display for the sake of visual assessment the restoration of 5th plane of an anisotropic 3D stack composed 18 images ^34^. The reconstruction quality is visibly superior with CARE as expected. Nevertheless SPITFIRE(e) with 3D Gaussian PSF (*σ*_*xy*_ = 2.0 pixels, *σ*_*z*_ = 0.5 pixel) and Gibson-Lanni model ^35^ (truncated to 3 planes around the center of the PSF) was able to provide satisfactory results. The main difference are visible on small filaments better enhanced with CARE. When we plot an intensity profile (**b**), we can see that it is the SPITFIR(e) method that gives the best compromise between pick thickness and rebound effect around the contours. The results can be visually improved with SPITFIR(e) if the volume is denoised beforehand and deconvolved with the PbyP strategy (Supplementary Fig. 8 (**a**), right).

## Discussion

An important challenge in live fluorescence imaging is to limit photodamages on the observed sample by reducing the dose of excitation light, while at the same time collecting enough photons to produce informative images for quantitative analysis. To overcome the difficulties of acquisition of 2D+Time and 3D+Time images at low excitation dose and boost the signal corrupted by blur and different sources of noise (shot noise, readout noise), we developed SPITFIR(e), a supermanoeuvrable and easy-to-use solution able to generate spatially consistent and high-quality fluorescence images. The approach is based on the fast minimization of a convex energy composed a novel sparse-promoting regularization term and a Poisson-Gaussian-aware quadratic fidelity term. Unlike previous methods which were tested on a few microscopy modalities ^13;14^ and need a careful adjustment of regularization parameters to obtain high-quality restoration results, SPITFIR(e) is a nearly automatized method which adapts to any microscopy technique, and image dimensionality. With this unique algorithm, the practitioner can switch from 4D image denoising to 3D deconvolution, or can chain the two processing tasks to obtain highquality images. He only needs to specify the level of desired sparsity (“low”, “moderate”, “high”) and to supply the measured PSF or the parameters of the 2D or 3D theoretical PSF (e.g., Gaussian) model.

We demonstrated on experimental data that SPITFIR(e) extracted high frequency information and achieved significant improvement in the output quality even with a poor calibrated PSF. Nevertheless, SPITFIR(e) could miss some details in noisy images taken at low excitation light intensity or short exposure time. As the lower light intensities approaches the limits of ability of SPITFIR(e) to recover information reliably, a successful strategy with consists in chaining denoising (3D or 4D) and deconvolution with a 2D or 3D PSF model to make the labeled biomolecules more discernible from noise. In presence of Poisson shot noise and readout noise, our results demonstrated that the proposed strategy is robust and able to boost signals reliably, and confirms that the quadratic fidelity term is suitable for Poisson-Gaussian noise as theoretically proved (Methods).

We have showed that SPITFIR(e) substantially reduces noise and provides better high-quality images than other currently available methods over a wide variety of experimental conditions. In the particular case of LLSM, which is designed already for long lasting fast imaging of 3D large and close to isotropic volumes, SPITFIR(e) shows all its power, since it not only improves resolution and reduces noise, but still allows an increase in the imaging rates by permitting to reduce the illumination dose. SPITFIR(e) also achieved high resolution in sub-diffracted and superresolution microscopy under low-light conditions, without perturbing the signal amplitudes. For two-dimensional live-cell STED imaging, line intensity profiles along mitochondria cristae showed that SPITFIR(e) reached an improved lateral resolution of order 1.5. Little is known about the actual dynamics of mitochondrial cristae in the multiple physiological processes to which mitochondria activity is required. This was mainly due to the visualization challenge of cristae structure in living cells in the past, both due to the lack of optical resolution and to the deleterious effect of light on mitochondria. Hessian-SIM modality ^36^ or STED nanoscopy ^37;38^ are the only approaches so far, able to achieve this goal. While only unique 2D STED images of live mitochondria are shown here, SPITFIR(e) applied on STED images might be of strong interest in this context since it allows to restore these fine structures even when depletion is incomplete and scanning imaging at very low illumination, making live relatively long term imaging accessible.

More generally, unlike smoothness regularizers, our sparse-promoting regularizer is able to reveal small details in the raw data, while preserving signal peaks and removing background, with no discernible artifacts in the final restored images. We also demonstrated that on 2D/3D benchmarks that SPITFIRE(e) outperforms existing deconvolution algorithms. Performance has been quantitatively studied in terms of PSNR values on artificially degraded 2D images and in terms of FRC on 3D experimental data. Finally, we showed that SPITFIR(e) is able to produce comparable results to supervised machine-learning techniques ^34^. Unlike content-dependent methods based on deep-learning algorithms requiring many training images, SPITFIR(e) is an unsupervised method which utilizes sparsity and smoothness as priori knowledge and adapts to any image contents.

In conclusion, our results on experimental data endorse that SPITFIR(e) may be considered as a “swiss-knife”, able to handle different samples captured under various fluorescence microscopes, adapt to various sources of signal degradation, image contents, and variable signal-to-noise ratios. This algorithm does not require a well-calibrated PSF and is easily controlled by choosing three levels of desired sparsity, that is “high”, “medium”, and “low”. We are convinced that SPITFIR(e) will be very helpful to push the spatiotemporal resolution limit of sub-diffracted and superresolution fluorescence microscopy techniques and will better resolve 3D structures and dynamics of biological components observed at low excitation light intensity to reduce artifacts in live cells. Our method was implemented as an Napari plugin (https://www.napari-hub.org/plugins/napari-sdeconv) for ease of use. It is also available as open source C++ code (CPU and GPU).

## Methods

### Variational model for image restoration

An observed 2D/3D image *f* : Ω ⊂ ℝ^*d*^ → ℝ, *d* = 2, 3 is a blurred and noisy version of the underlying true image *u* : Ω → ℝ (i.e., *u* ∈ ℝ^Ω^). A general approach for restoring an image consists in finding the minimizer of an energy functional, i.e.,

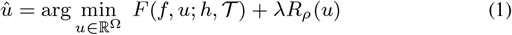

where *F*(*f, u*; *h, 𝒯*) and *R*_*ρ*_(*u*) are the data fidelity term and the regularization term, respectively, and *λ >* 0 and *ρ* ∈ [0, 1] are the parameters controlling the amount of smoothness and sparsity in the image *u*. The fidelity data term is the distance between the restored image *u* and the observed image *f* where *h* is the PSF function and 𝒯 is the degradation operator that encompasses Poisson and Gaussian noise. In what follows, we give the explicit expressions of the two terms.

### Data fidelity term

A fidelity data term is generally derived from the general formation model, for instance dedicated to low photon counts and low-light regimes in fluorescence microscopy ^13^. In our approach, *u* ∈ Ω is assumed to be corrupted by a mixed Poisson-Gaussian. The local noise variance is then represented at a given location **x** ∈ Ω as follows ^39;40;60^ (see details in Supplementary Notes (Appendix A)):

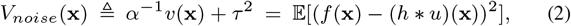

where *f* is the noisy image, *v* = *h* * *u*, 𝔼[·] denotes the mathematical expectation, *α >* 0 is the quantization factor of the photodetector, and *τ*^2^ *>* 0 represents the Gaussian noise variance. If we assume that the average intensities in *v* and *f* are preserved, it follows that:

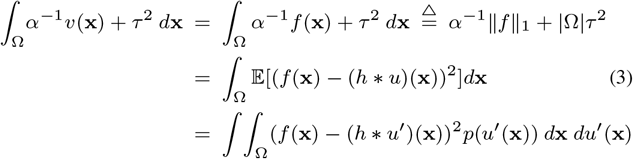

where *p*(*u′* (**x**)) represents the probability distribution of *u*. If we assume the following prior *p*(*u′* (**x**)) = **1**[*f*(**x**) = ((*h* * *u*)(**x**))] where **1**[·] is the indicator function, we finally obtain:

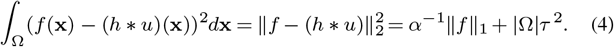

Starting from the seminal paper^41^, the restored image is found by solving the following optimization problem (Poisson-Gaussian noise):

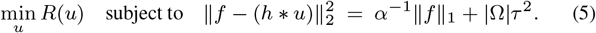

It turns out that (5) is a constrained formulation with an equality constraint which is not convex. The corresponding formulation with inequality constraint is

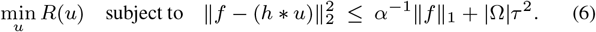

It has been established that, under additional assumptions, that (5) and (6) are equivalent. A Lagrange formulation can be then derived since the the right-hand side of the inequality does not depend on the unknown image *u*:

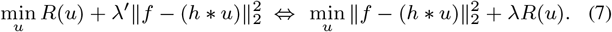

where the parameter *λ′ >* 0 balances the two energy terms with *λ′* = *λ*^−1^ ≠ 0 (see Supplementary Notes (Appendix A) for details). In conclusion, a quadratic fidelity term *F*(·) is appropriate for Gaussian-noise removal provided it is combined with regularizer *R*(·).

### Regularization term

Commonly-used regularizers in image restoration have the following form:

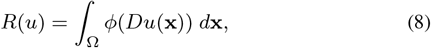

where *D* is a linear operator (called “regularization operator”) used to control the spatial distribution of *u* and *ϕ*(·) is a positive potential function usually related to a norm distance. A typical example is the Tikhonov-Miller regularization ^8^ defined as the squared norm of the gradient of *u*: 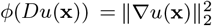.

We propose a convex sparsity-promoting regularizer based on second-order derivatives to avoid the emergence of undesirable stair-casing effects. The Sparse Hessian Variation (SHV) regularizer is defined as follows

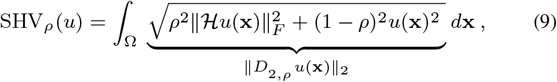

where ‖ · ‖_*F*_ denotes the matrix Frobenius norm and *ρ* ∈ [0, 1] is the weighting parameter. Moreover, since the Frobenius norm of the Hessian matrix is equal to the Euclidean norm of its vectorized version, the operator *D*_2,*ρ*_ ∈ ℝ^10^ is defined (for *d* = 3) as:

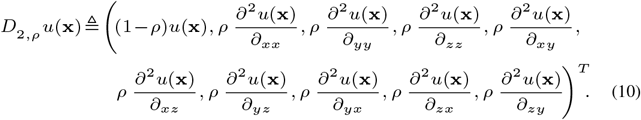

The idea behind this combination is to sparsify jointly the spatial distribution of image intensities and the second order derivatives of the image to encourage smooth variations between spatially-contiguous non-zero regions of the underlying image. The resulting image contain bright objects against a large smooth dark background, and no spurious edge. Unlike LSHV ^14^, SHV is convex and more flexible as it involves a parameter *ρ* ∈ [0, 1] which is helpful to eliminate or preserve background. In SPITFIR(e), we consider only three values to control the amount of sparsity: *ρ* = 0.1 (“sparse”), *ρ* = 0.6 (“moderately sparse”), and *ρ* = 0.9 (“weakly sparse”).

### Discrete formulation and optimization

We present here the discretization setting required to derive the optimization algorithm. Let us consider a sampling grid of the following form in 3D (*d* = 3) (expressions in 2D are similar).

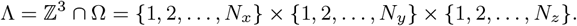

The observed noisy and blurry image *f* is represented by its digitized (discrete) version as follows: *f* = 𝒯 (*Hu*), where 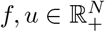 with *N* = *N*_*x*_ ×*N*_*y*_ ×*N*_*z*_, *H* ∈ ℝ^*N* ×*N*^ is a matrix that models the point spread function of the microscope in the discrete setting, and 𝒯 is degradation operator. Both *u* and *f* are assumed to be non negative. In the discrete setting, the blurring operator *H* corresponds to a discrete convolution which can be efficiently computed by using fast Fourier transform (FFT) ^42;43;44;45^. To discretize *D*_2,*ρ*_, we use standard finite differences for the Hessian operators with Neumann conditions on image boundaries. The 2D-3D deconvolution problem is then defined in the discrete setting as the minimizer of the following energy:

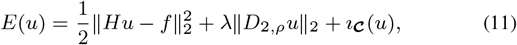

where *λ >* 0 is the regularization parameter and *ı*_***𝒸***_ is the characteristic function of the convex subset 𝒸 defined as: *ı*_***𝒸***_ = 0 if *u* ∈ *𝒸* and +∞ otherwise, is very helpful to impose positivity constraint on the solution.

The optimization problem (11) is convex since the underlying energy functional is defined as the sum of convex terms, but it is non-smooth. To minimize the energy (11), several algorithms have been investigated, including FISTA ^46^, ADMM ^22;23^ and MM ^47^ algorithms. Instead of applying the aforementioned optimization methods, we investigated a first-order method to minimize the sum of convex functions, based on the proximal splitting approaches ^25;26;48;49;50;51;52;53^. The main idea consists in splitting the original problem into several simple sub-problems in the way that each single function of the sum can be processed separately (see details in Supplementary 1C). These operators are well-suited for large-scale problems arising in signal and image processing, because they only exploit first-order information of the function and thus enable fast and efficient computation. To solve the minimization problem (11), we adapted the full splitting approach described in ^25;26^. The key idea is to evaluate the gradient, proximity and linear operators individually in order to avoid implicit operations such as inner loops or inverse of linear operators.

### Self-tuning of the regularization parameter

The adjustment of the regularization parameter *λ* in (11) is generally crucial in most image restoration algorithms, and may be time consuming. A non optimal choice of this parameter may over-smooth object borders, suppress structural details, generate artifacts or weakly reduce noise. Therefore, we investigated a self-tuning strategy which is based on the comparison of restorations and performance criteria over a range of parameters *λ*. The practical issue is to automatically tune this parameter on a case-by-case basis to get the best performance as possible, from the input noisy image. To address this issue, we propose a self-tuning approach based on the analysis of one or several informative regions of interest (ROI) in the input noisy image. The ROI can be the entire image or a *p*×*p* patch containing significant fluorescent signal and randomly drawn or manually selected by the user in the input image.

Instead of applying the principle of cross-validation, we investigated here an approach based on the minimax principle which avoids the selection of low and high values for the regularization parameter ^54^. First the energy (1) is rewritten as follows:

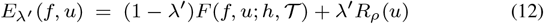

where *λ′* ∈ ]0, 1[ is a positive constant such that *λ′* = *λ*(1 + *λ*)^−1^. The MinMax principle amounts to finding *u*^⋆^ that minimizes, over all *λ′* values, the maximum of *E*_*λ*′_(*f, u*):

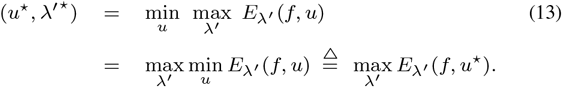

The optimization problem may be solved by examining the finite set of solutions *E*_*λ*′_(*f, u*^⋆^) and selecting the *λ′* valued associated to the highest energy *E*_*λ*′_(*f, u*^⋆^). The good news is that Gennert and Yuille ^54^ demonstrated that there exists a unique value *λ′* that maximizes *E*_*λ*′_(*f, u*^⋆^) (convexity of *E*_*λ*′_(*f, u*^⋆^). Hence, by initializing *λ′* = 0.5 and considering a finite set 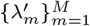 of increasing values such that 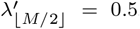, a fast search technique can be developed which consists in restoring the image with 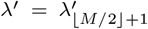 and 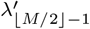 and keeping the *λ*′ corresponding to the highest energy *E*_*λ*′_(*f, u*^⋆^). The raw image is then restored with a highest (or lowest value) of *λ′*, and the SPITFIR(e) procedure is repeatedly applied on raw data (with *λ′* = *λ′* (1 − *λ*^*′*^)^−1^) until the energy *E*_*λ*′_(*f, u*^⋆^) decreases (or increases).

In practice, the level of sparsity (“sparse”, “moderately sparse”, “weakly sparse”) is first selected by the practitioner beforehand. The parameter *λ′* is then estimated by applying the aforementioned optimization procedure to a single or multiple 2D ROIs (or patches) located in informative areas, that is in areas containing meaningful fluorescent signals, as illustrated in Supplementary Fig. 9.

### Implementation details and code availability

Images have been processed on classical workstation for CPU calculation (CPU Intel Xeon 2.8GHz 4 threads). We use this setup since it is the most common setup in biology labs. For high speed processing we developed a GPU version of SPITFIR(e). We run the GPU on two types of GPU, a Nvidia Quadro M2000 that can be found in most workstations and a Nvidia Tesla K80 for computing grid. Typical computing times (CPU and GPU) for processing 2D-3D-4D images are reported in the table below: These computing times are small and take into account the full data processing steps (including data normalization and data copy to the GPU device). The code can be downloaded for free from the GitLab website (https://github.com/sylvainprigent/simglib) along with accompanying documentation.

### Data source

See description in Appendix C (Microsopy and Cell Cultures), Supplementary Information.

## Acknowledgements

This work was jointly supported by the French National Research Agency (France-BioImaging ANR-10-INBS-04-07, DALLISH DALLISH-ANR-16-CE23-0005), and INNOPSYS company. We would like to thank Mathieu Morin from Inserm U932 for his help with STED image acquisition, and the PICT imaging platform of Institut Curie, member of the France-BioImaging infrastructure (ANR-10-INBS-04-01) for maintaining spinning disk and LSM confocals used in this study.

## Author contributions

C.K. devised the project and the main conceptual ideas, supervised the project and was in charge of overall direction and planning. C.K., H.-N.N. and S.P. designed and implemented the presented SPITFIR(e) method, in discussions with J.S., C.A. V.-C. and L.L. J.S. C.A. V.-C. and L.L. provided the real datasets (LLSM, STED, CM) datasets and conceived experiments and implemented the analysis workflows applied to real images. B.H. provided the real MFM datasets. H.-N.N. and S.P. performed experiments on artificial datasets and real images (WF, MFM, LLSM, CM). C.K., H.-N.N. and S.P. co-wrote the manuscript. All authors provided critical feedback and helped shape the research, analysis and manuscript.

## SUPPLEMENTARY INFORMATION

### SUPPLEMENTARY VIDEOS

**Suppementary Movie 1**. Composite movie corresponding to the time series of insets in Fig. 2 **a, b, e** and **f**. Upper part from left to right, microtubules, mitochondria and superimposed MIP data from LLSM after deskew treatment. Lower part, similar disposition after SPITFIR(e) 4D denoising and 3D deconvolution. Only 1:45 minutes of a 4 minutes series is shown. Scale bar: 5 *μ*m.

**Supplementary Movie 2**. Multi-Angle view of the 3D of superimposed image stacks from the same time series as in Supplementary Movie 1. Upper part data from LLSM after deskew treatment, lower part after SPITFIR(e) 4D denoising and 3D deconvolution. Scale Bar = 5 *μ*m.

### SUPPLEMENTARY FIGURES

**Supplementary Fig. 1:**
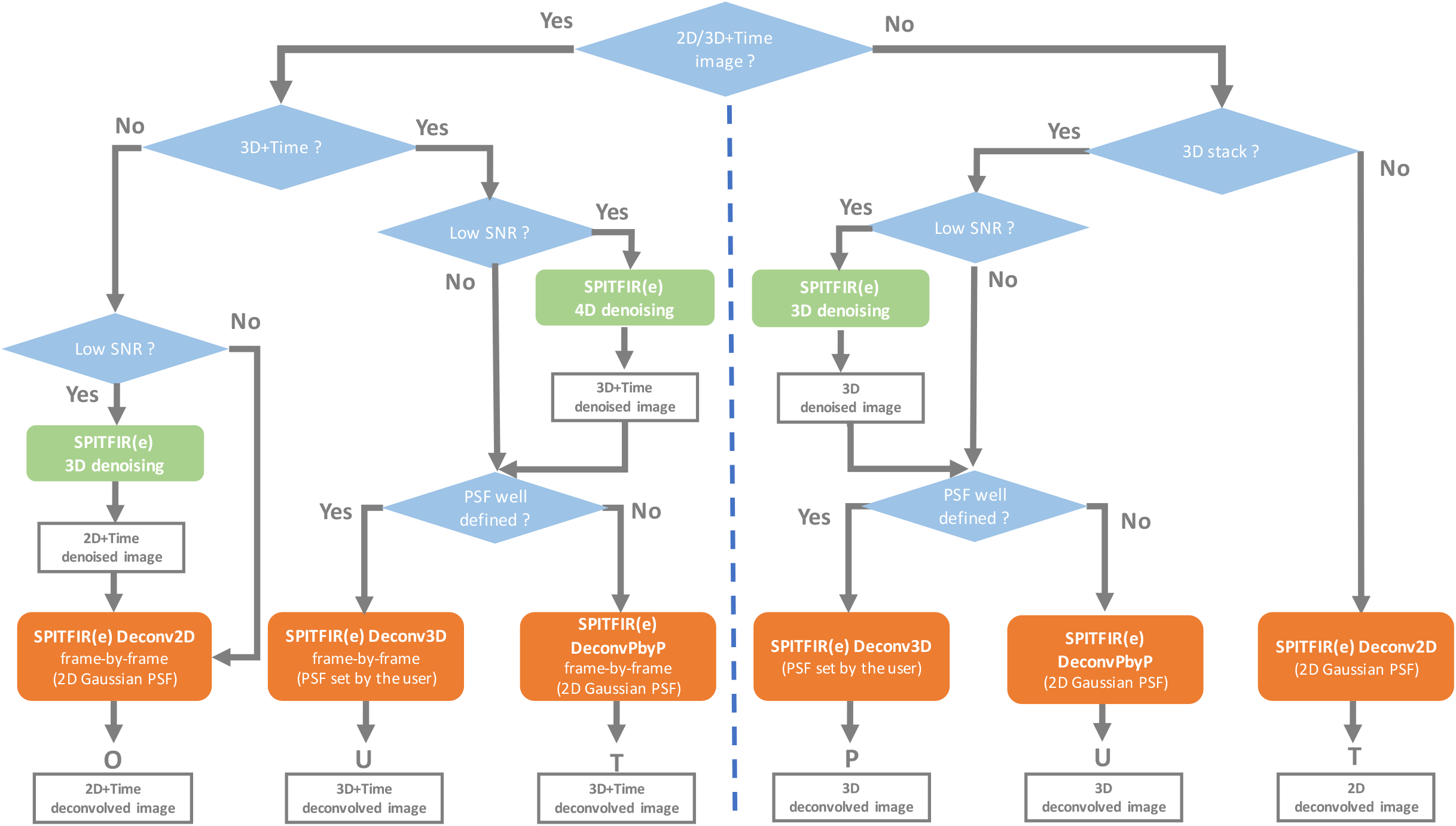
SPITFIR(e) overview. The flow chart describes the sequence of restoration steps applied to 3D, 2D+Time, or 3D+Time images based on responses of end-users.

**Supplementary Fig. 2:**
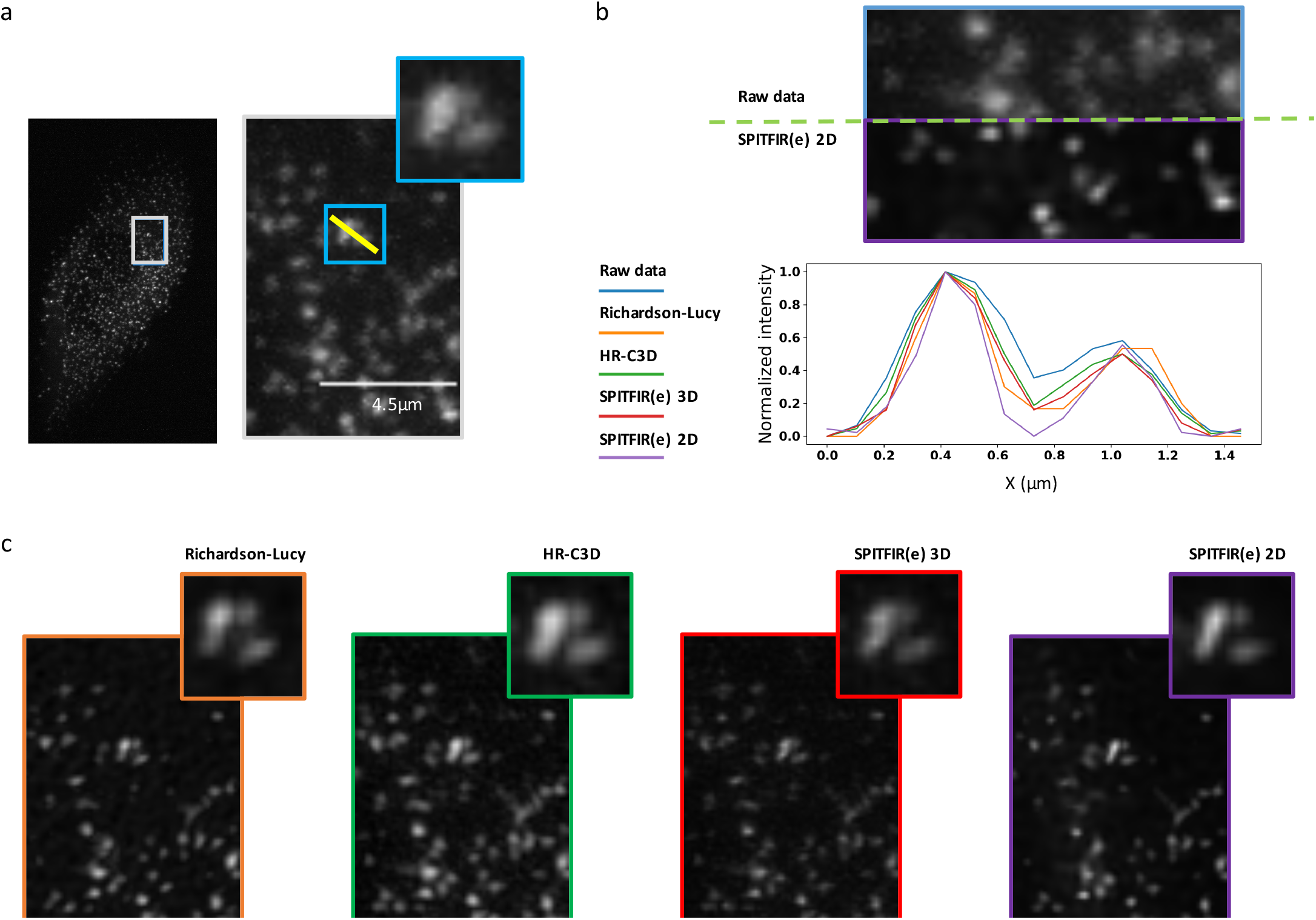
Deconvolution of a 3D LLSM image single plane image depicting *σ* subunit-eGFP of the AP2 complex (scale bar: 4.5 *μ*m, source: CNRS UMR144 Institut Curie) with SPITFIR(e) and HR-C3D. (**a**) Original image (raw data) extracted from a 3D volume of 58 planes. The pixel size is 104 nm along the XY directions and 325 nm along the z axis. (**b**) The intensity profiles for the four methods (Richardson-Lucy, HR-C3D, SPITFIR(e) 3D and SPITFIR(e) 2D) are measured along the yellow segment superimposed within a ROI (red window) in (**a**). (**c**) Images obtained with the four different deconvolution algorithms. SPITFIR(e) 3D used a 3D Gaussian PSF model with parameters *σ*_*xy*_ = 1.0 pixel and *σ*_*z*_ = 0.8 pixel. SPITFIR(e) 2D used a 2D Gaussian PSF with parameter *σ*_*xy*_ = 1.0 pixel. Scale bar: 4.5 *μ*m.

**Supplementary Fig. 3:**
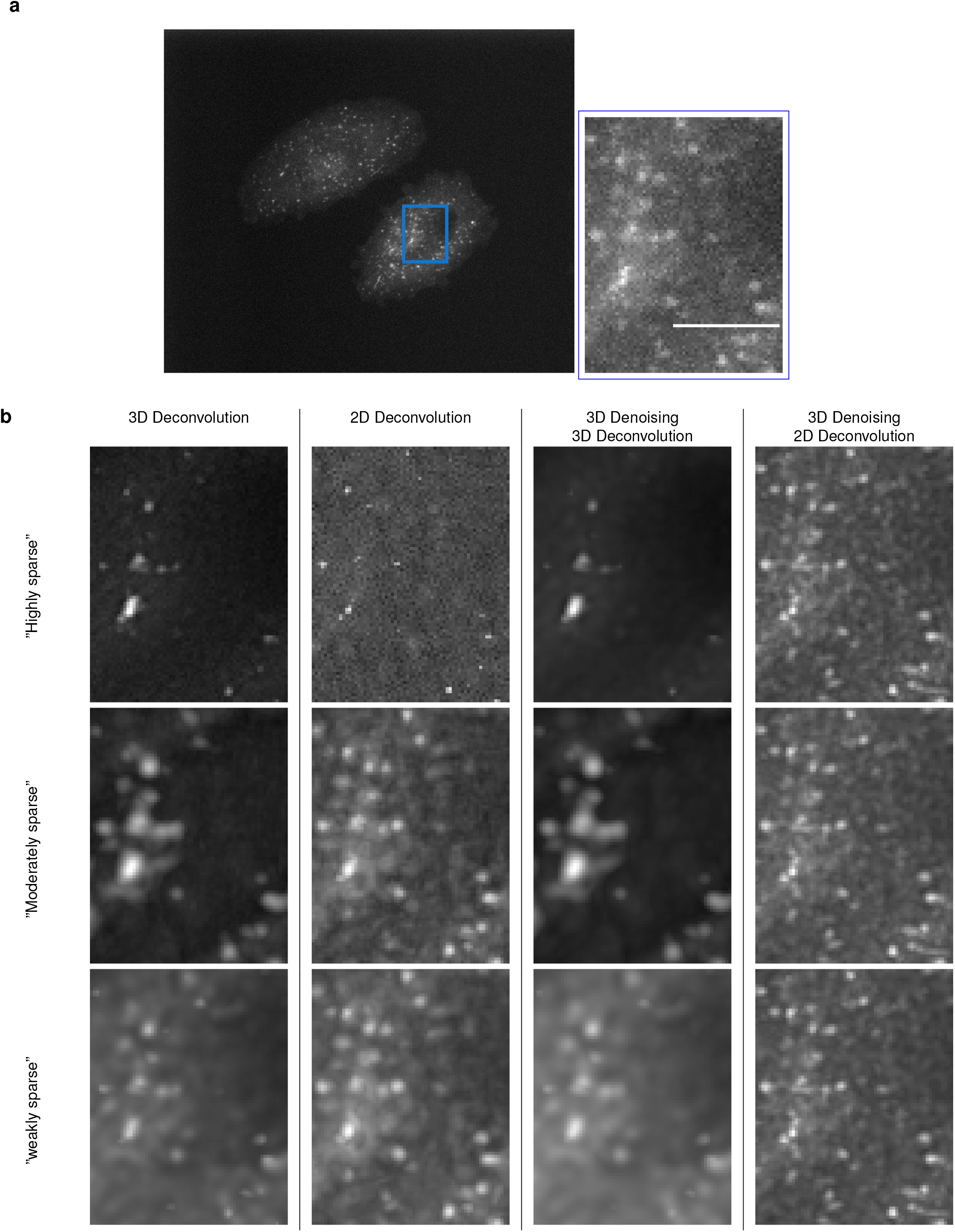
Deconvolution of 3D spinning disk confocal microscopy image. **a**, Display of a 2D plane depicting Hela cells expressing GFP-Rab5 proteins (scale bar: 4 *μ*m). **b**, Results with SPITFIR(e) with different amounts of sparsity (“weak” (*ρ* = 0.9), “moderate (*ρ* = 0.6), “high” (*ρ* = 0.1), automatic selection of the regularization parameter, and different strategies with SPITFIR(e): 2D Deconvolution, 3D Decovolution, 3D Denoising + 3D Deconvolution, 3D Denoising + 2D deconvolution (*σ*_*xy*_ = 1.5 pixels and *σ*_*z*_ = 0.5 pixel).

**Supplementary Fig. 4:**
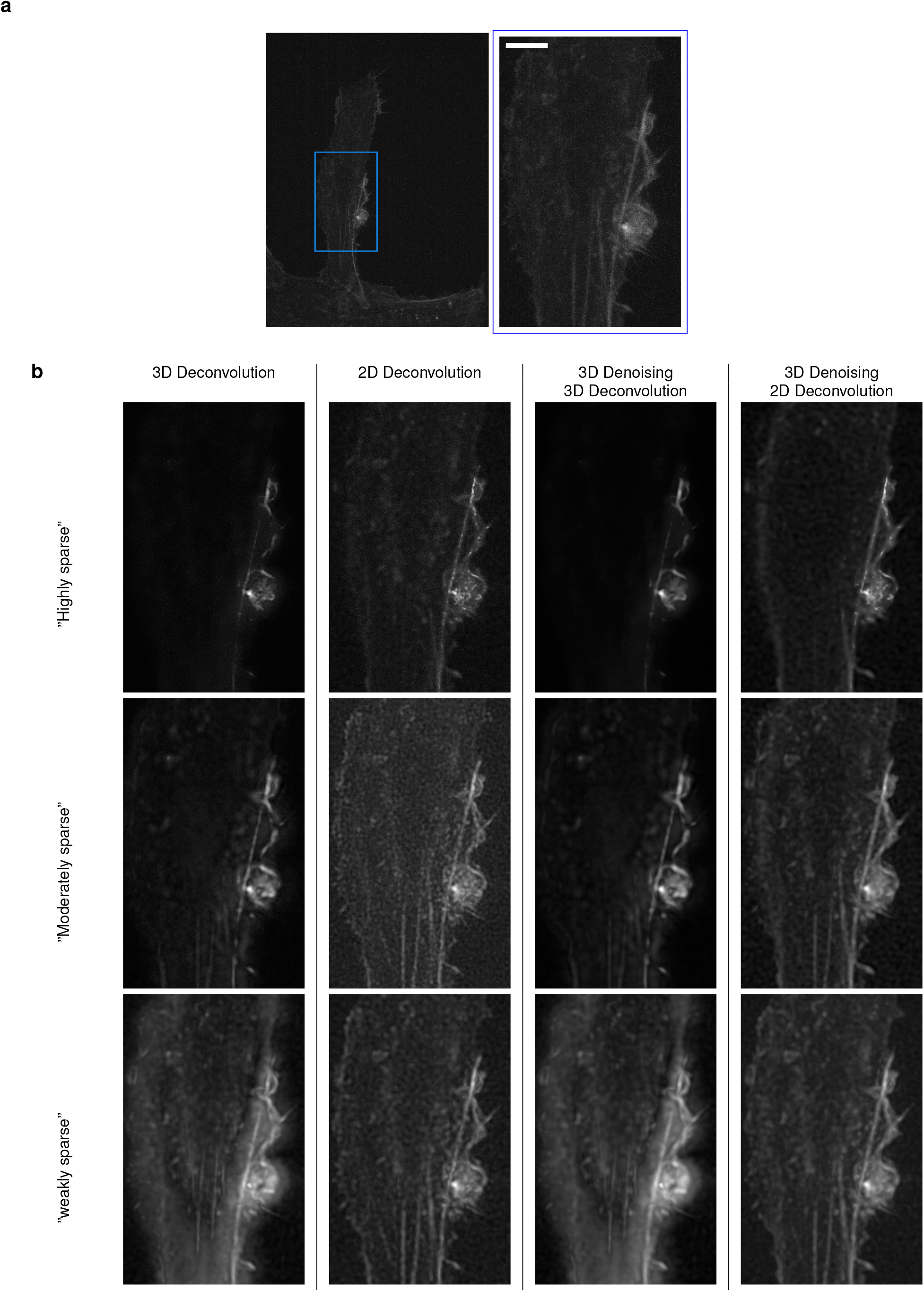
Deconvolution of 3D spinning disk confocal microscopy image. **a**, Image depicting mCherry-LifeAct in RPE1 cells (scale bar: 5 *μ*m). **b**, Results with SPITFIR(e) with different levels of sparsity (“weak” (*ρ* = 0.9), “moderate (*ρ* = 0.6), “high” (*ρ* = 0.1), automatic selection of the regularization parameter and different strategies with SPITFIR(e): 2D Deconvolution, 3D Decovolution, 3D Denoising + 3D Deconvolution, 3D Denoising + 2D deconvolution (*σ*_*xy*_ = 1.5 pixels and *σ*_*z*_ = 0.5 pixel).

**Supplementary Fig. 5:**
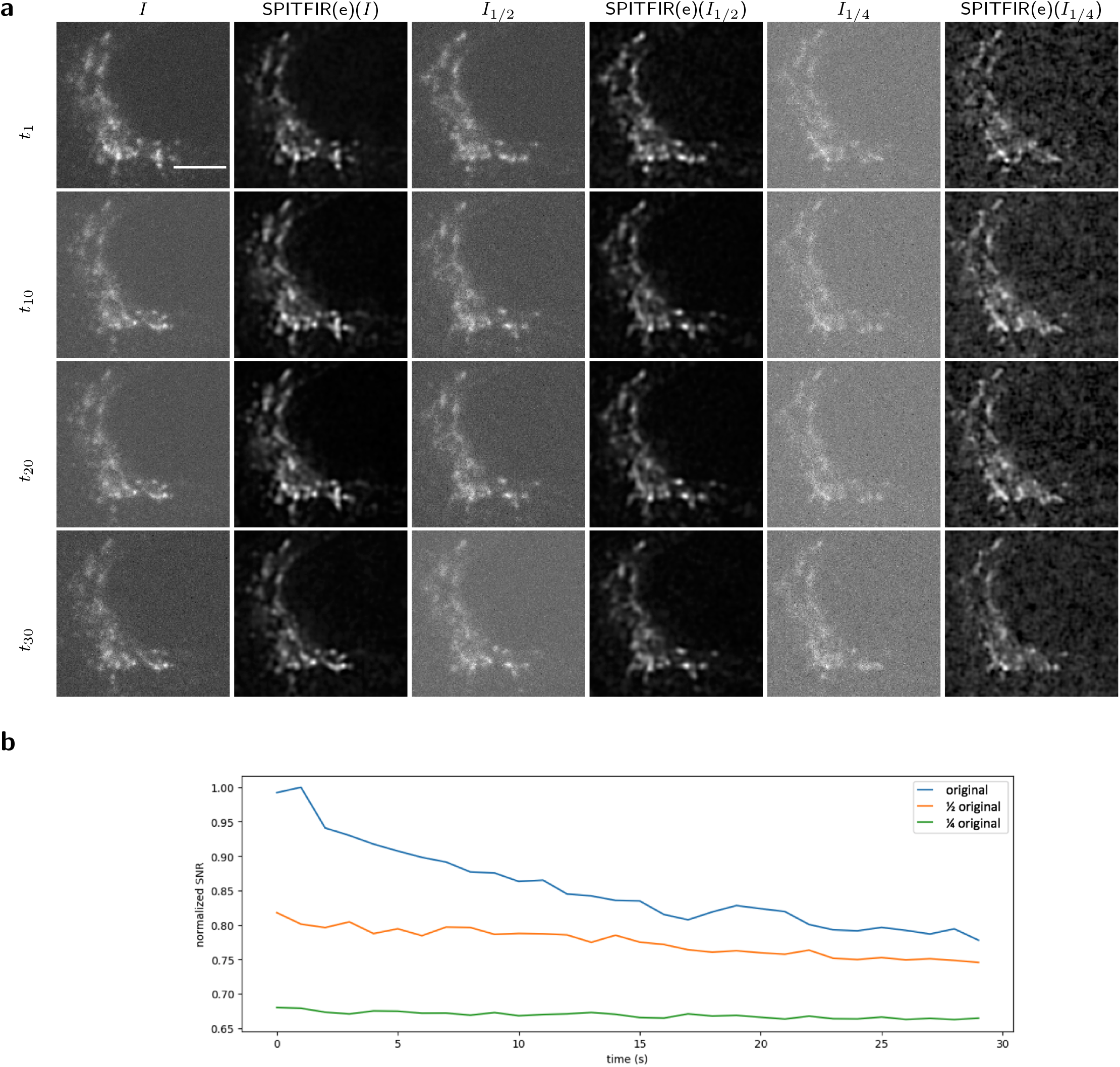
SPITFIR(e) (4D denoising + 3D deconvolution) applied to spinning disk confocal microscopy images. **a**, Data are temporal series (30 time points) of 3D stacks composed 14 planes each depicting CD-M6PR-eGFP Hela cells. The sample is imaged 3 times with different doses of illumination light. The figure depicts the 6th plane extracted from the 3D stack of a region of interest (ROI) at time *t*_1_, *t*_10_, *t*_20_, and *t*_30_. Scale bar = 4.5 *μ*m. The *I* image is illuminated with high dose of illumination light to get an image with a high SNR. The *I*_1*/*2_ image is illuminated with half power compared to *I*. The *I*_1*/*4_ image is obtained by dividing the dose of illumination light by a factor 4 compared to *I*. We applied SPITFIR(e) (4D denoising + 3D deconvolution with a Gibson-Lanni PSF model truncated to 3 planes around the center of the PSF and the acquisition parameters) to the three 4D volumes. The results are denoted SPITFIR(e)(*I*), SPITFIR(e)(*I*_1*/*2_) and SPITFIR(e)(*I*_1*/*4_), respectively. **b**, The plot shows the SNR of the three original movies over time.

**Supplementary Table 2:**
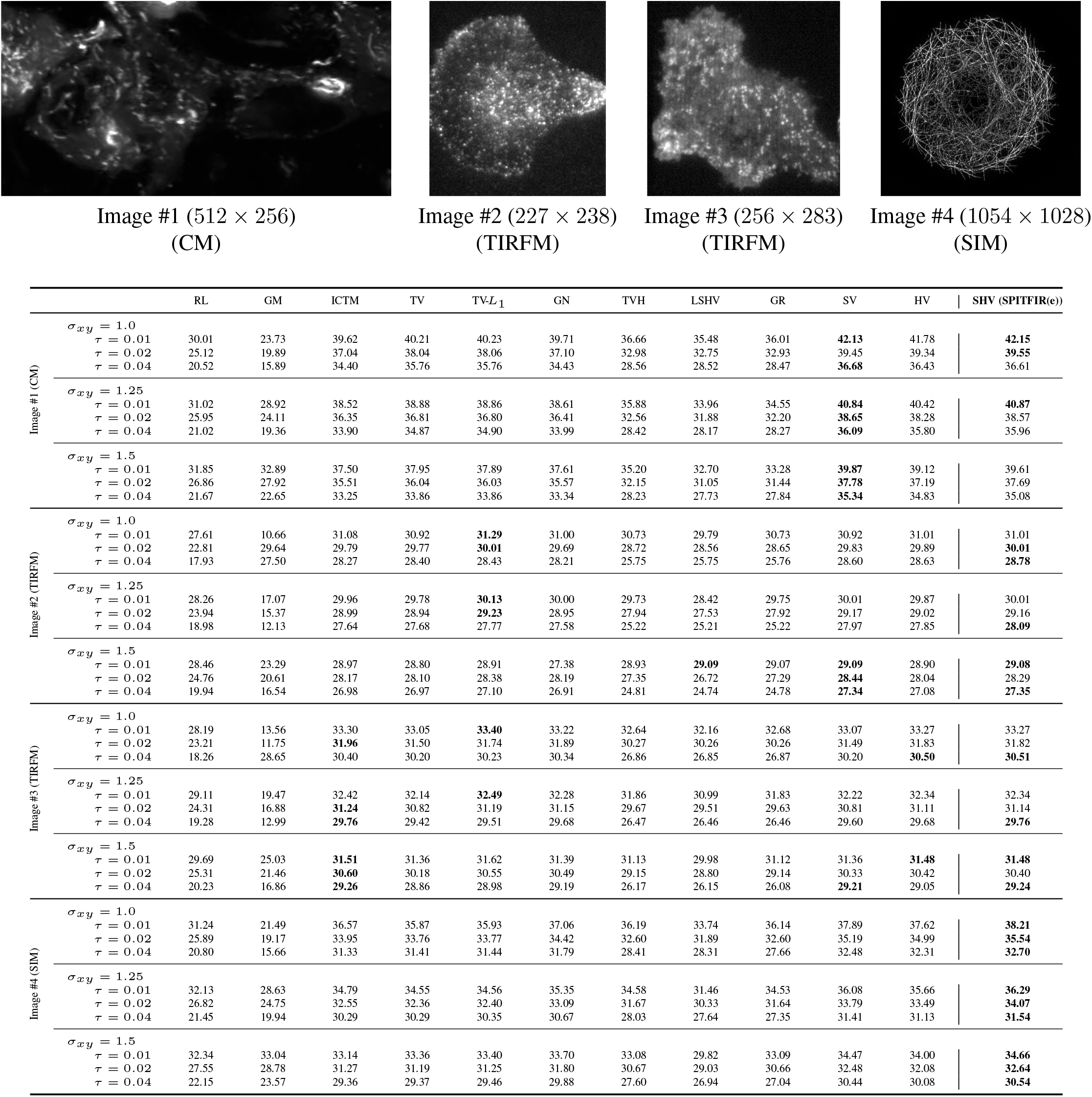
PSNR scores of deconvolution methods applied to four 2D images (confocal (CM), TIRFM, SIM). The best scores (±0.05 dB) are in bold style. Images #1-#3 depict fluorescently tagged proteins (Rab11-mCherry, Langerin-YFP, TfnR-pHluorin (Transferin-receptor)) corresponding to small bright spots over a dark background, observed in spinning confocal microscopy (CM) (Image #1, courtesy of PICT-IBiSA imaging platform) and total internal reflection fluorescence microscopy (TIRFM) (Images #2 and #3) respectively, previously acquired ^55;56^. Image #4 depicts microtubules observed in Structured Illumination Microscopy (SIM) with high resolution (up to 100 nm) This image (CIL 36147) is taken from the Cell Image Library (CIL) (http://www.cellimagelibrary.org). The images were deconcolved with the following methods: Richardson-Lucy (RL) ^3^ algorithm, Gold-Meinel (GM) ^57^ algorithm, iterative constrained Tikhonov-Miller (ICTM) ^7^ algorithm, Total Variation (TV) ^41^ regularizer, Total-Variation-L1 (TV-L1) ^19^, GraphNet (GN) ^15^ regularizer, Total Variation-Huber (TVH) ^58^ regularizer, Log Sparse Hessian Variation (LSHV) ^14^ regularizer, Good’s roughness (GR) ^59^ regularizer, Sparse Variation (SV) ^18^ regularizer, Hessian Variation (HV) ^12^ regularizer, and Sparse Hessian Variation (SHV) (our SPITFIR(e) method) regularizer.

**Supplementary Fig. 6:**
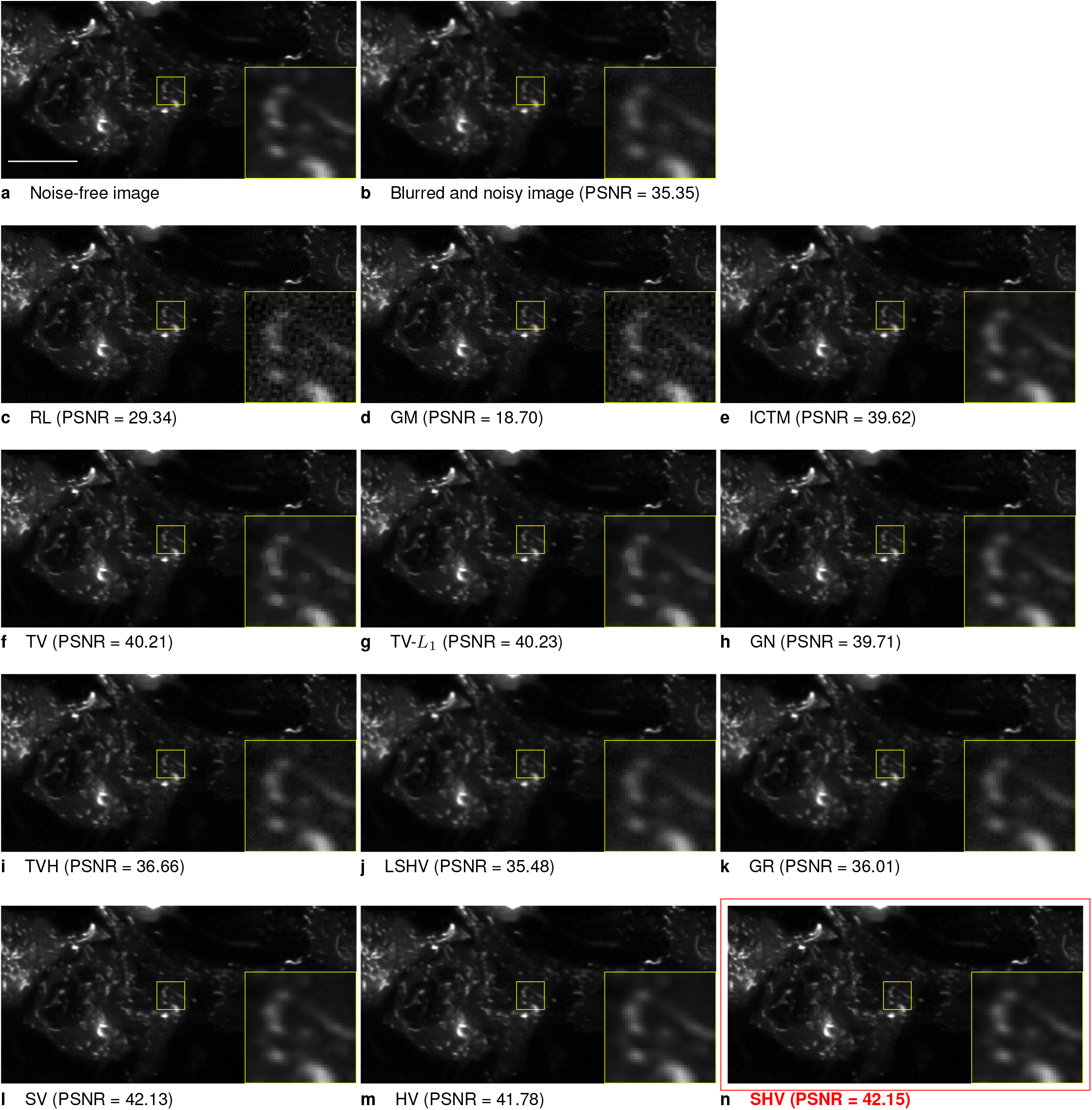
Deconvolution of a confocal microscopy image with different methods. **a**, The original 512 × 256 pixels image (Image #1) (source: CNRS UMR144 Institut Curie) depicting Rab11a-mCherry protein in M10 cells, was previously acquired in confocal spinning-disk microscopy. Scale bar: 10 *μ*m. **b** Artificially corrupted image by a Gaussian PSF *σ*_*xy*_ = 1.0 pixel and Gaussian white noise with standard deviation *τ* = 0.01 (**b**). **c**, image deconvolved with the Richardson-Lucy (RL) ^3^ algorithm. **d**, image deconvolved with the Gold-Meinel (GM) ^57^ algorithm. **e**, image deconvolved with the iterative constrained Tikhonov-Miller (ICTM) ^7^ algorithm. **f**, image deconvolved with the Total Variation (TV) ^41^ regularizer. **g**, image deconvolved with the Total-Variation-L1 (TV-L1) ^19^. **h**, image deconvolved with the GraphNet (GN) ^15^ regularizer. **i**, image deconvolved with the Total Variation-Huber (TVH) ^58^ regularizer. **j**, image deconvolved with the Log Sparse Hessian Variation (LSHV) ^14^ regularizer. **k**, image deconvolved with the Good’s roughness (GR) ^59^ regularizer. **l**, image deconvolved with the Sparse Variation (SV) ^18^ regulariser. **m**, image deconvolved with the Hessian Variation (HV) ^12^ regularizer. **n**, image deconvolved with the Sparse Hessian Variation (SHV) (our SPITFIR(e) method) regularizer (red box). Zoom-in views are displayed in yellow windows for comparison in details.

**Supplementary Fig. 7:**
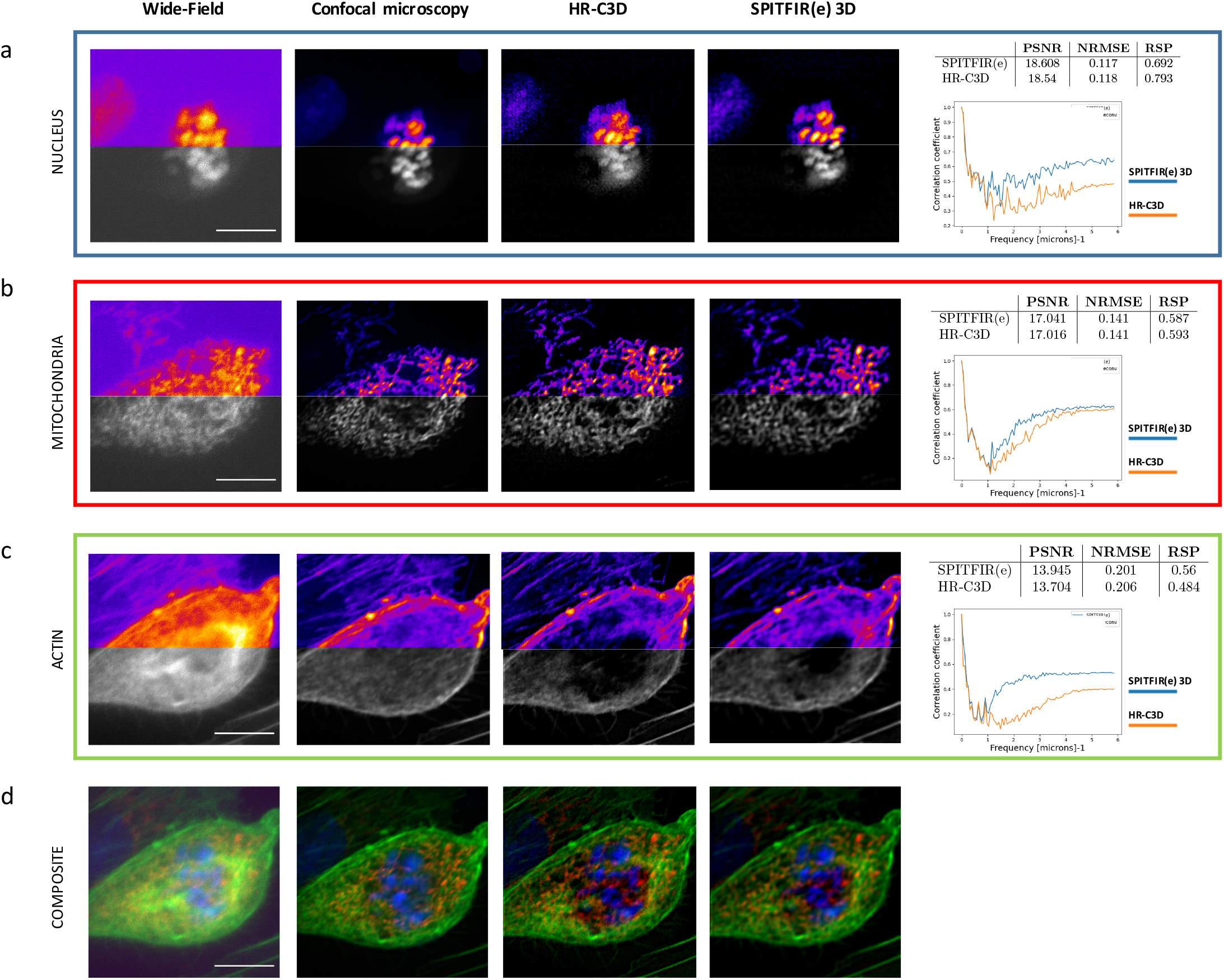
A comparison of SPITFIR(e) 3D and HR-C3D ^13^ applied to a 3D wide-field microscopy image. Display of 2D (512 × 512 pixels) images extracted from a stack composed of 22 planes depicting nucleus stained with DAPI (**a**, blue channel, 11th plane), mitochondria staind with MitoTracker Red CMXRos (**b**, red channel, 6th plane), and actin staind with lexa Fluor 488 phalloidin (**c**, green channel, 4th plane) (see description in ^13^). **d**, Display of the wide-field composite image (11th plane, merging of the three channels), the corresponding confocal image deconvolved with Huygens software (see ^13^) which will serve as high-resolution reference image, and the deconvolved images with HR-C3D and SPITFIR(e) 3D, from left to right, respectively. The three plots show the Fourier Shell Correlation (FSC) between the wide-field image deconvolved with the two competing methods and the reference (spinning disk) confocal image. Each plot corresponds to one color channel. The Peak-Signal-to-Noise-Ratio (PSNR), the Normalized Root Mean Square Error (NRMSE) and the global Pearson. correlation coefficient (RSP) are reported in Tables. Scale bar: 10 *μ*m.

**Supplementary Fig. 8:**
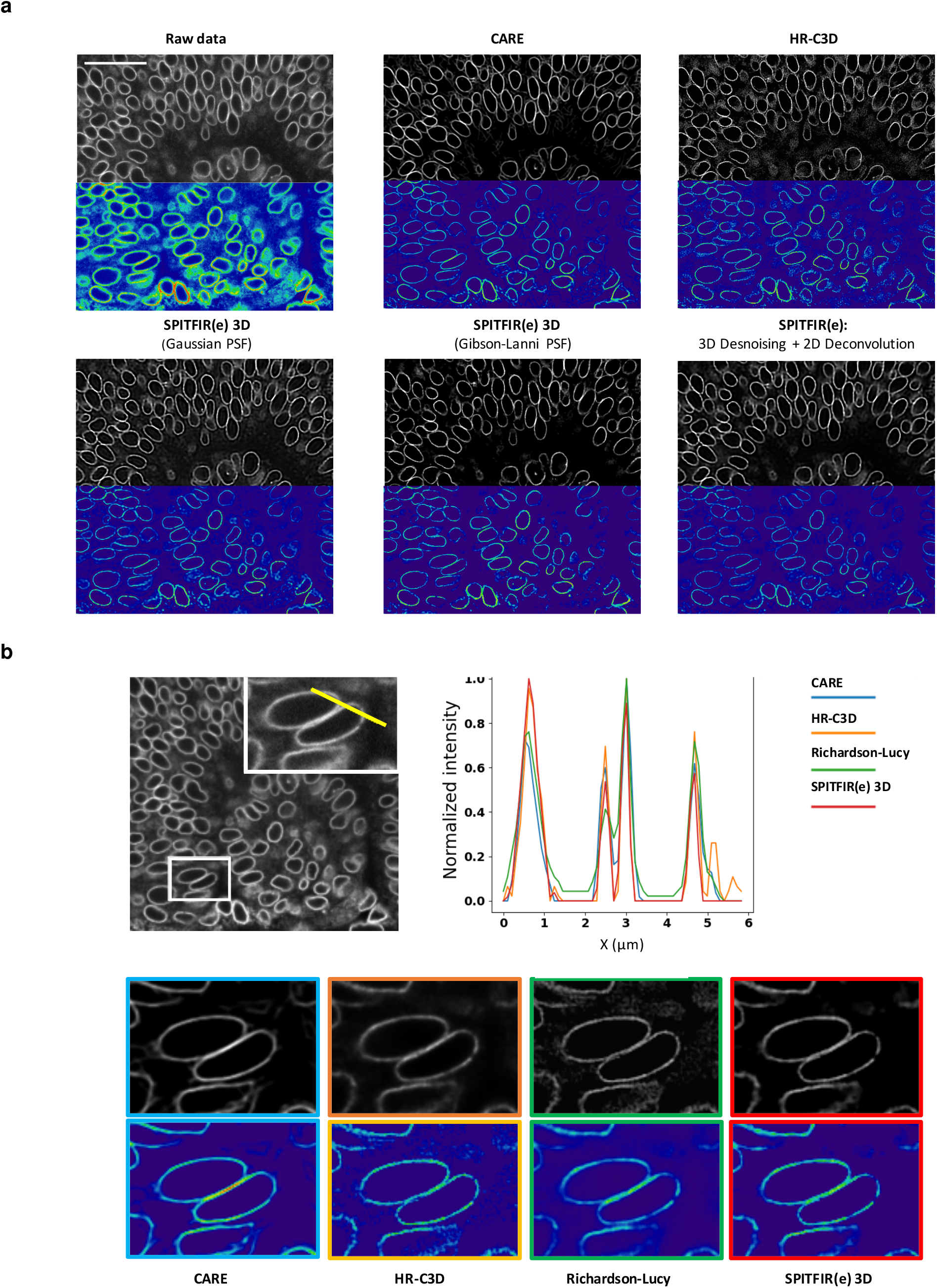
Deconvolution of a 3D spinning disk confocal microscopy image depicting the envelopes of nuclei stained with GFP-LAP2b (developing eye of zebrafish (Daniorerio) embryos, source: ^34^) (scale bar: 25 *μ*m). **a**, Comparison of RL, HR-C3D, CARE and SPITFIR(e) with a Gibson-Lanni PSF modeld and a 3D Gaussian PSF model. We display the results for the 5th plane. **b**, The plots show the intensity profile for each 3D deconvolution method along the line drawn in yellow in ROI (white rectangle). Bottom, Zoom-in views (5th plane) of deconvolution results obtained with CARE, RL, HR-C3D and SPITFIR(e) 3D (Gibson-Lanni (generated from ^34^: 60X/1.3-NA objective, 488 nm, z step = 2 *μ*m) or 3D Gaussian PSF model (*σ*_*xy*_ = 2.0 pixels and *σ*_*z*_ = 0.5 pixel).

**Supplementary Fig. 9:**
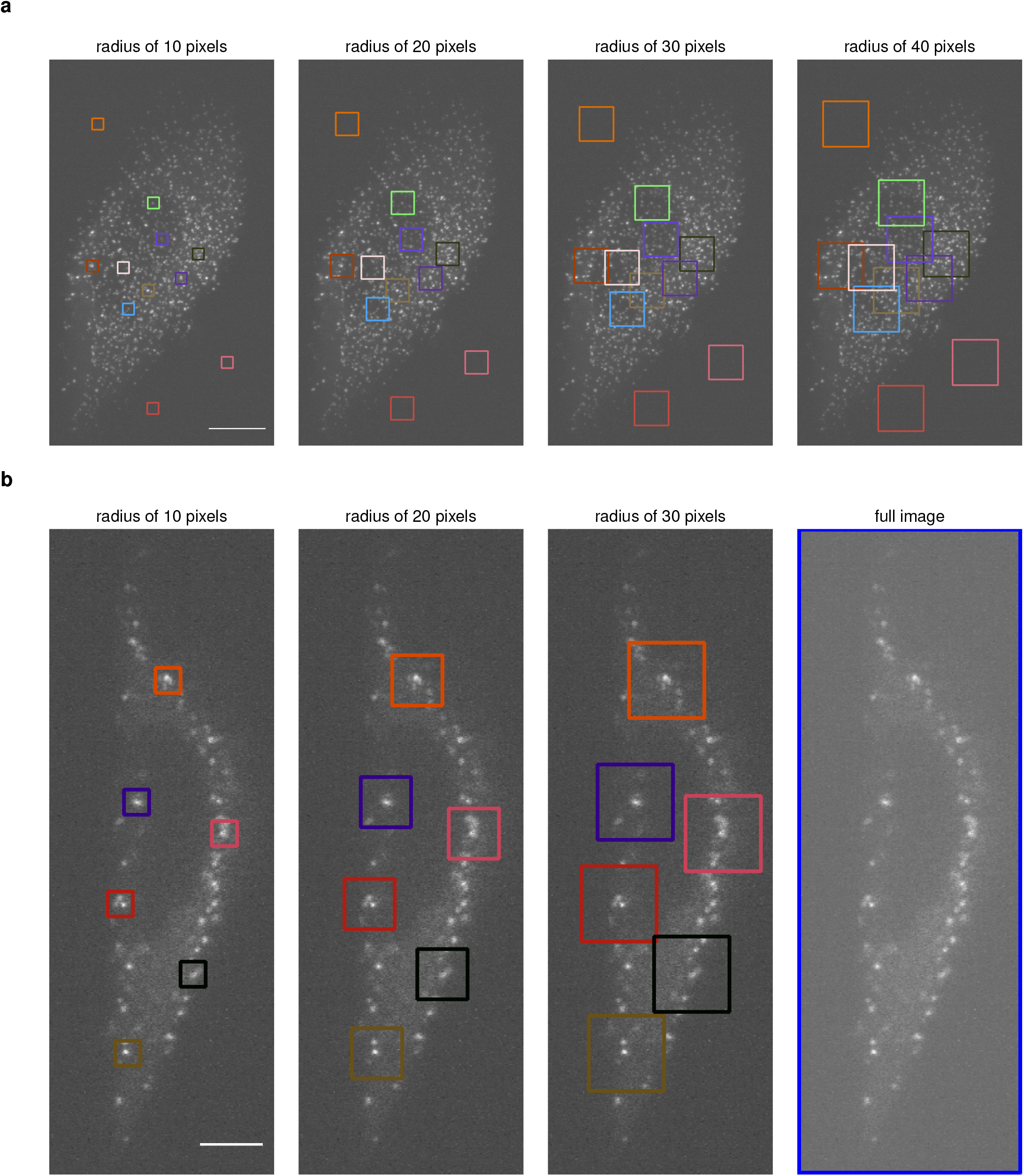
Automatic selection of *λ* parameter using the minimax principle applied to ROIs with variable radius. **a**, Display of ROIs with radius of 10, 20, 30 and 40 pixels superimposed on the maximum projection image of 3D LLSM image depicting *σ* subunit-eGFP of the AP2 complex in Hela cells. Scale bar: 10 *μ*m. **b**, Display of ROIs with radius of 10, 20, and 30 pixels superimposed on the maximum projection image of 3D LLSM image depicting *σ* subunit-eGFP of the AP2 complex in Hela cells (Orientation XY based on Fig. 1). Scale bar: 5 *μ*m.

### SUPPLEMENTARY NOTES

#### A. Definition of the energy for images corrupted by Poisson-Gaussian noise

A fidelity data term is generally derived from the general formation model, for instance dedicated to low photon counts and low-light regimes in fluorescence microscopy (e.g., ^13^). We formally demonstrate below that a conventional quadratic fidelity term, which is optimal when the images are corrupted by additive white Gaussian noise, is also appropriate for mixed Poisson-Gaussian noise.

##### Calculation of noise variance

Let us consider the following observation model ^60^:

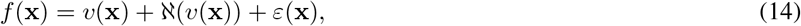

with *v*(**x**) = (*h* * *u*)(**x**), where **x** ∈ Ω is the pixel position in the domain Ω, ℵ and *ε* represent the signal-dependent Poisson noise component and the zero-mean Gaussian noise component respectively, such that:

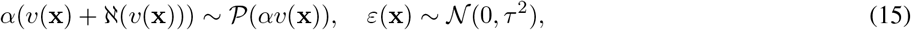

*α >* 0 is the quantization factor of the photodetector, and *τ*^2^ *>* 0 represents the Gaussian noise variance. According to the properties of Poisson distribution (𝔼[·] and Var[·] denote the mathematical expectation and the variance respectively), we have

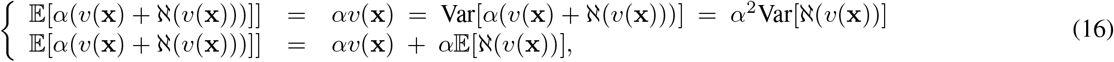

which yields to 𝔼[ℵ(*v*(**x**))] = 0 and Var[ℵ(*v*(**x**))] = *v*(**x**)*/α*. Hence,

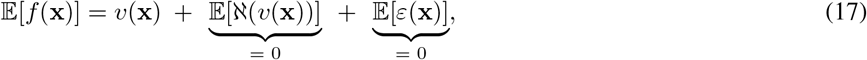

and the overall variance of *f*(**x**) is the sum of Var[ℵ(*v*(**x**))] and Var[*ε*(**x**)] = *τ*^2^:

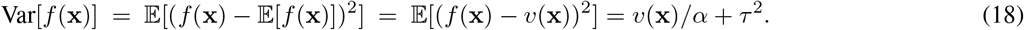

We conclude that *v*(**x**) is defined as the expected value of the noisy observations *f*(**x**) and Var[*f*(**x**)] = *α*^−1^*v*(**x**) + *τ*^2^ is the noise variance ^39;40;60^.

##### Definition of energy model

Assume an image *v* = *h* * *u* corrupted by a mixed Poisson-Gaussian and denote *f* the noisy image. As explained above, the local noise variance can be represented at a given location **x** ∈ Ω as:

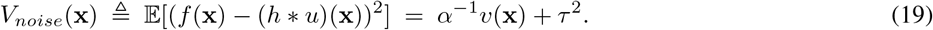

Let us assume that the average intensity is preserved, that is

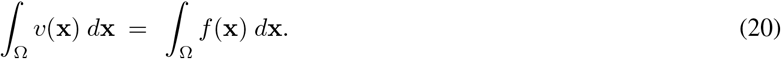

It follows that, by pre-multiplying the intensities on both sides of the previous equation by a factor *α*^−1^ and adding the constant value *τ*^2^, we have:

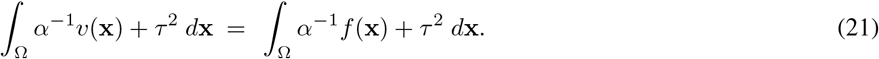

Meanwhile, by integrating (19) over the image domain Ω, we get

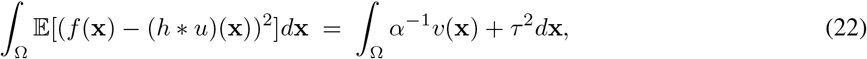

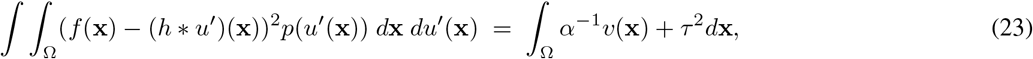

where *p*(*u′* (**x**)) represents the probability distribution of *u*. Assume the following prior *p*(*u′* (**x**)) = **1**[*f*(**x**) = ((*h* * *u*)(**x**))] where **1**[·] is the indicator function. Hence, we have by using (21)

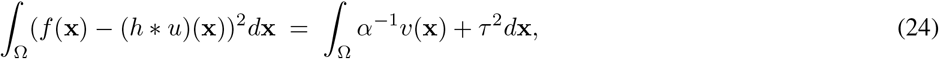

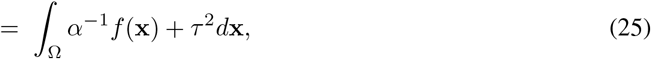

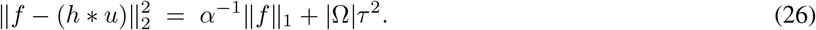

Now, starting from the seminal paper^41^, the restored image is found by solving the following optimization problem (Poisson-Gaussian noise):

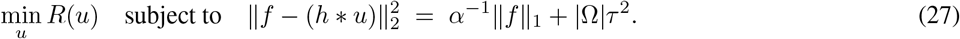

It turns out that (27) is a constrained formulation with an equality constraint which is not convex. The corresponding formulation with inequality constraint is

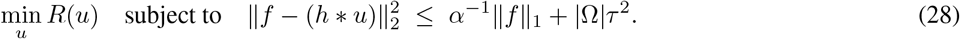

It has been established that, under additional assumptions, that (**??**) and (28) are equivalent. A Lagrange formulation can be then derived since the the right-hand side of the inequality does not depend on the unknown image *u*:

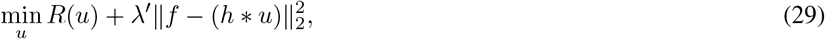

where the parameter *λ′ >* 0 balances the two energy terms and is an unknown function of the Poisson-Gaussian noise variance. The Karush-Tucker conditions guarantees that (30) and (28) are equivalent for a particular choice of *λ′*. Finally, if we set *λ′* = *λ*^−1^ ≠ 0, we can equivalently reformulate the minimization problem as:

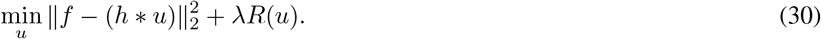

#### B. Discrete formulation and optimization

##### Discrete formulation

The observed noisy and blurry image *f* is represented by its digitized (discrete) version and not by its continuously defined counterpart. The continuous model is not appropriate for discrete images even though the estimation of the continuous image *u* from discrete samples of *f* is in principle possible. Consider a discrete formulation by assuming that the images *u* and *f* are non negative and sampled according to the sampling grid

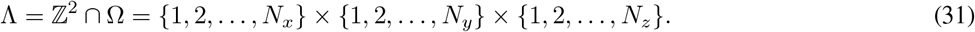

The observed noisy and blurry image *f* is represented by its digitized (discrete) version as follows:

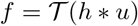

where 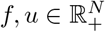 with *N* = *N*_*x*_ × *N*_*y*_ × *N*_*z*_, *H* ∈ ℝ^*N* ×*N*^ is a matrix that models the point spread function of the microscope in the discrete setting, and 𝒯 is the degradation operator. For a coordinate (*i, j, k*) ∈ Λ, we denote by *u*_*i,j,k*_ (resp. *f*_*i,j,k*_) the value of *u* (resp. *f*) at position (*i, j, k*) ∈ Λ. A discrete version of these images are therefore given by 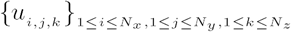 and 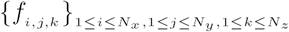. Denote *χ* = ℝ^*N*^ with *N* = *N*_*x*_ × *N*_*y*_ × *N*_*z*_, a finite dimensional vector space equipped with a standard inner (scalar) product

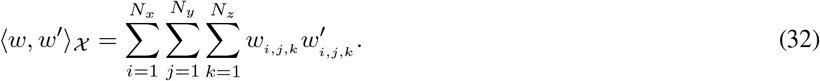

The induced norm by the defined inner product is given by

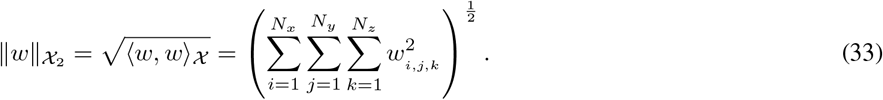

In the discrete setting, the blurring operator *H* corresponds to a discrete convolution which can be efficiently computed by using fast Fourier transform (FFT) ^42;43;44;45^. To discretize *D*_2,*ρ*_, we use standard finite differences to approximate the second derivatives along the three dimensions of the volume at voxel (*i, j, k*), with Neumann conditions on image boundaries:

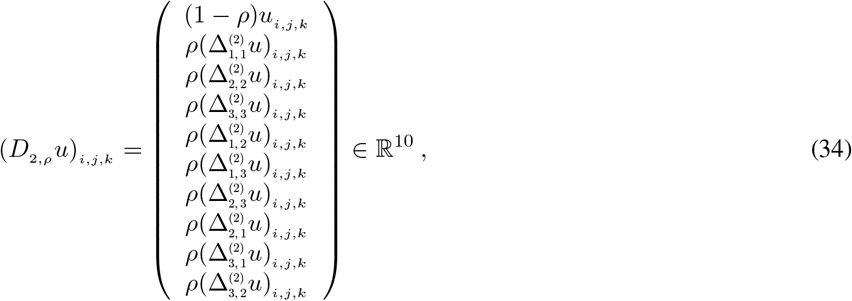

where

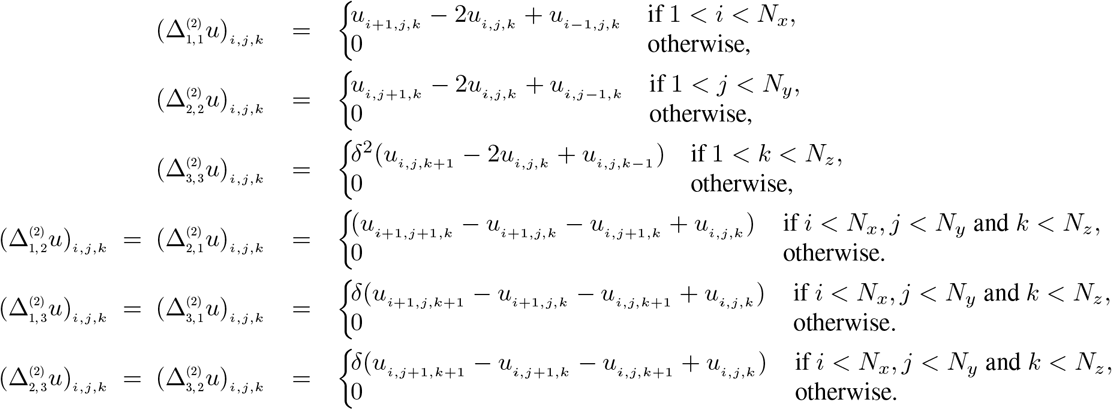

and *δ* is the ratio of the lateral-to-axial step sizes.

The discrete operators can be used to define the discrete SHV regularizer as:

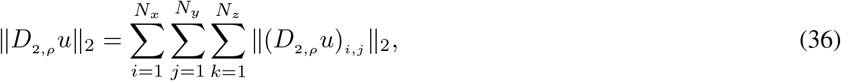

where the *L*_1_-norm acts now on the discrete domain Λ. The 3D deconvolution problem is defined in the discrete setting as the minimizer of the following energy:

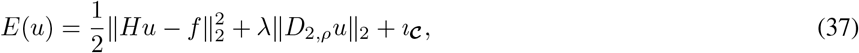

where *λ >* 0 is the regularization parameter and *ı*_***𝒸***_ is the indicator of a convex set 𝒞 such as:

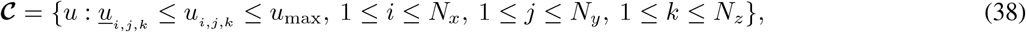

where the upper bound *u*_max_ *>* 0 is the maximal intensity value allowed and *u*_*i,j,k*_ ≥ 0 is an estimated lower bound of the pixel intensity which is spatially varying and then adapted to each pixel location. The spatially varying constraint on the lower bound of pixel intensity not only guarantees positivity of the solution but also helps to avoid over-sparsifying effect. In our experiment, for the sake of simplicity, we use the following lower bound 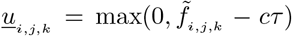 where *c >* 0 and 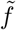 is a smoothed version of the observed noisy image *f* by a low-pass (Gaussian) filter.

##### Energy minimization and splitting algorithms

We notice that the objective function

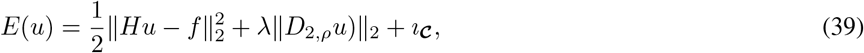

is a sum of linear composite functions as 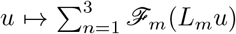, where each *ℱ*_*m*_ is a convex function and each *L*_*m*_ is a linear bounded operator. Formally, we can write *ℱ*_1_ = *ı*_***𝒸***_, *L*_1_ = Id, *ℱ*_2_ = *λ* ‖ ·‖_2_, *L*_2_ = *D*_2,*ρ*_ and 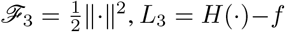. Generic primal-dual proximal approaches can be used to minimize this linear combination of convex functions as proposed in^61;62^, but it is not optimal since the smoothness of the quadratic terms 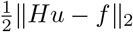 is not exploited. In order to solve the problem (39), the design of an appropriate algorithm requires therefore to take into account the specific form of the corresponding energy, i.e., the sum of a simple convex function *ℱ* = *ı*_***𝒸***_, a more sophisticated composite function *𝒢* ∘ *L* = *λ* ‖ *D*_2,*ρ*_ (·)‖_2_ (here, *𝒢* = *λ* ‖ · ‖_2_ and *L* = *D*_2,*ρ*_) and a smooth function 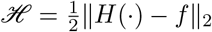.

In what follows, we present a first-order method to minimize the sum of convex functions based on the proximal splitting approaches ^25;26;48;49;50;51;52;53^. It consists in decomposing (splitting) the original problem into several simple sub-problems in the way that each single function of the sum can be processed separately. Indeed, smooth function involves its gradient operator, while non-smooth function implies its Moreau proximity operator ^63^. These operators are well-suited for large-scale problems arising in signal and image processing, because they only exploit first-order information of the function and thus enable fast and efficient computation.

Let us recall first that the proximity operator of a convex function *𝒥* : ℝ^*N*^ → ℝ is defined as

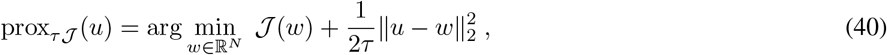

where *τ >* 0 is a control parameter. From this definition, it easy to verify that the proximity operator of the function *ℱ* (*u*) = *ı*_***𝒸***_ (*u*) is nothing else than the projection onto the convex subset ***𝒸*** as the following

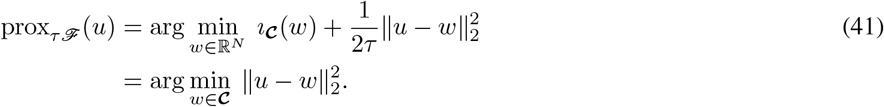

If we denote proj_***𝒸***_ the projection operator on ***𝒸***, its closed-form expression is given by

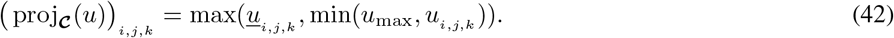

Moreover, the quadratic function 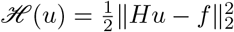 possesses an analytic form for its associated proximity operator

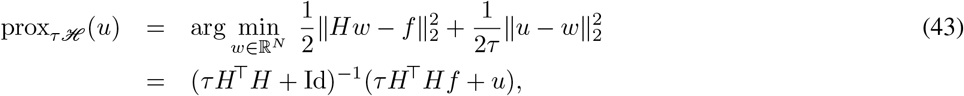

where the symbol ^T^ denotes the adjoint of a linear operator and *H*^T^ : ℝ^*N*^ → ℝ^*N*^ satisfies 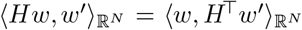. The evaluation of prox_*τ ℋ*_ (*u*) corresponds to the inverse of a linear system that is not always possible in practice due to the high dimensionality of the problem. For this reason, the optimization methods which involve the gradient of *ℋ* are more appropriate since they do not require any inverse operator. In the comparison with *ℱ* and *ℋ*, the calculation of the proximity operator in the case of the composite function *𝒢 ∘ L*(*u*) = *λ‖D*_2,*ρ*_ *u ‖*_2_ is theoretically possible but is challenging because of the presence of *D*_2,*ρ*_ which is not diagonal.

To solve the minimization problem (39), we adopt the full splitting approach described in ^25;26^. The key idea of this approach is to evaluate the gradient, proximity and linear operators individually in order to avoid implicit operations such as inner loops or inverse of linear operators. Accordingly, only “simple” computations are considered such as the gradient ∇*ℋ*, the proximity operator of *ℱ* and *𝒢*, the linear mapping *L* and its adjoint operators *L*^T^. The corresponding proximal algorithm for the problem is written under the following general form at iteration *𝓁*:

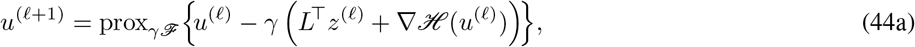

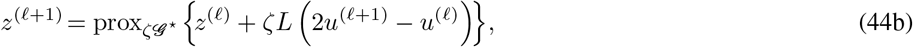

where *γ, ζ >* 0 are proximal parameters, *𝒢*^*\**^ denotes the Legendre-Fenchel conjugate of *𝒢* and its proximity operator 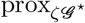 can be directly computed from 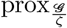 by using the Moreau’s identity 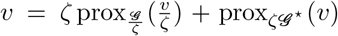. Following ^25;26^, to guarantee the convergence of the proposed algorithm, the parameters *γ* and *ζ* must fulfill the condition

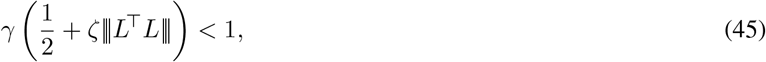

where ||| · ||| denotes the operator norm. The proofs of the convergence can be found in ^25^. We also note that the proposed algorithm belongs to the class of primal-dual algorithms which provide not only the solution of the primal problem (a.k.a. the original minimization problem) but also the solution of its dual problem.

Since the closed-form of prox_*γℱ*_ is already given, it remains to define the analytic expression of other terms in (44a) and (44b). We start with the gradient of the quadratic function *ℋ* which is straightforwardly obtained by

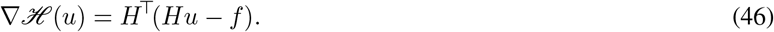

Next, we notice that the regularization operator *L* = *D*_2,*ρ*_ is a linear mapping, then its adjoint operator 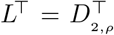 is defined using the equation 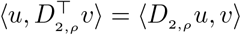, which implies (*d* = 3)

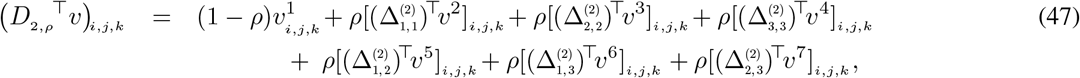

where the involving adjoint operators are given below:

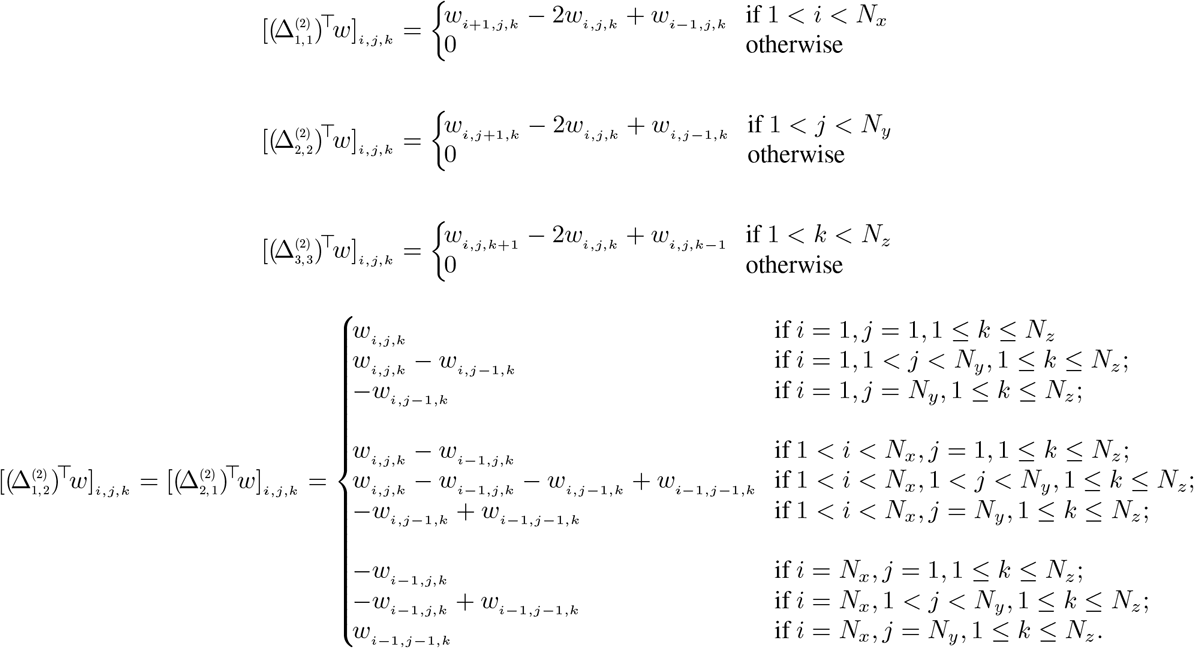

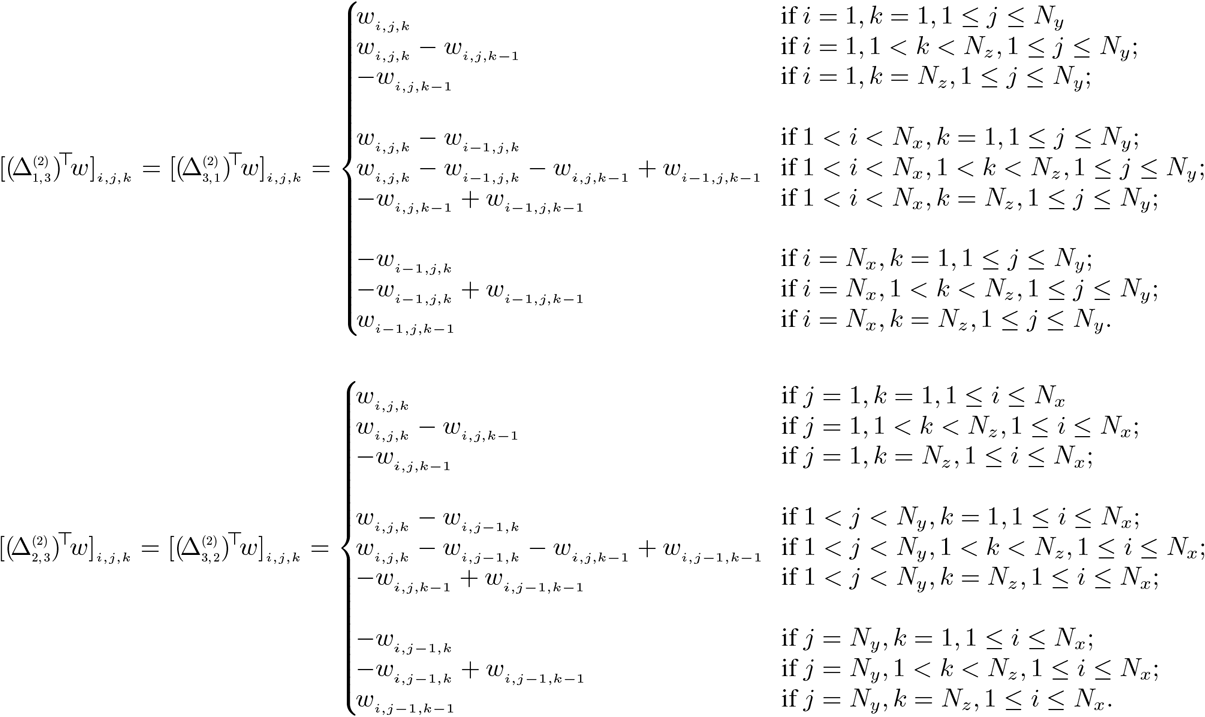

From equations (47), one can deduce the following upper bound:

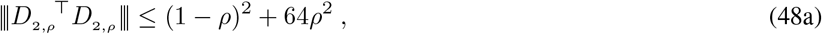

that are used for choosing the proximal parameters *γ* and *ζ* according to (45).

The last term we want to deal with is the proximity operator 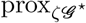. We also note that the proposed primal-dual algorithm does not necessitate evaluating the proximity operator of the composite function *𝒢* ∘ *L* as in the case of generic proximal algorithms, but only 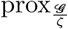 is required. Since *𝒢* is related to the mixed norm whose the proximity operator is defined as:

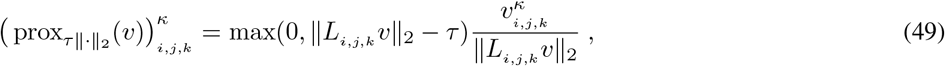

where 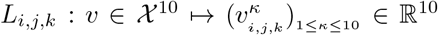 is a linear operator (*d* = 3). By using the Moreau’s identity, we obtain the closed-form expression of prox_*ζ𝒢*_^*\**^ as the following:

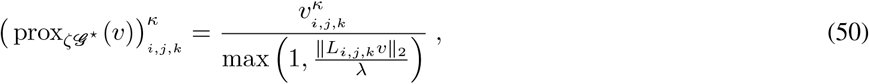

which shows that 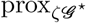 is independent from *ζ* and moreover it is an pointwise operator. These properties allow therefore fast and efficient computation by exploiting the intrinsic parallelism of multicore processors.

##### Energy model for 4D denoising

A temporal series of 3D noisy images *f* : Ω ⊂ ℝ^4^ → ℝ is a noisy version of the underlying true 3D image sequence *u* : Ω → ℝ modeled as follows: *f* = 𝒯 (*u*). It follows that the denoising problem is formulated as the minimization of an energy functional defined as

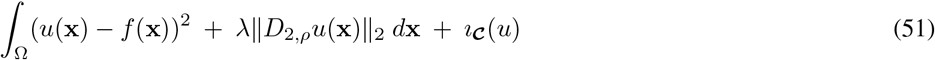

where

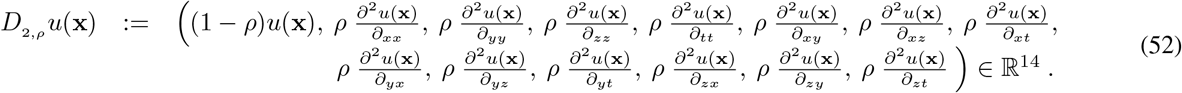

To solve the optimization problem, we split the original problem (51) into several simple sub-problems as explained above. The resulting individual functions involved in the sum are then minimized separately. The algorithm is expected to be faster since FFT is not required in the implementation for image denoising. Nevertheless, the amount of data is much more larger since the whole 4D image sequence is denoised at once. In practice, a very long image sequence can be segmented into sub-sequences with a small overlap, and is processed independently and in parallel. The overlapping images can be averaged to reduce possible artifacts at the end.

#### C. Microscopy and cell cultures

##### Cell culture fluorescence labeling

The hTERT-immortalized RPE1 cells (Human Retinal Pigment Epithelial Cell) were purchased from ATCC (CRL-4000). RPE1 cells within 30 passage number were used and grown in DMEM-F12 medium without phenol red (Life Technologies) supplemented with 10% (vol/vol) fetal bovine serum (FBS) at 37°C in humidified atmosphere with 5% CO2. For LLSM (Lattice Light Sheet Microscopy) imaging, cells were seeded 5 to 6 hours before image acquisition at 150.103 per well of a 6 well plate, containing 3 to 4 coverslips with a 5mm diameter, treated as previously described in (Chen *et al*. 2012)^1^. Live cells were stained for microtubule and mitochondria using the Tubulin TrackerTM DR (Tub-Tracker Deep Red, ThermoFisher Scientific) probe (Ex. 652nm/Em. 669) at 10 ng/ml and 100ng/ml of PKMR (PK Mito Red), a cyclooctatetraene-conjugated-cyanine-3 dye (Ex. 549/Em. 569) kindly provided by Dr. Z. Chen from Peking University (Yang *et al*. 2020)^2^, respectively. Cell labeling was performed in two steps. Cells were first incubated for 15min with Tubulin Tracker DR and washed twice with LLSM medium (DMEM/F-12, without phenol red, with 1% BSA, 0.01% penicillin–streptomycin, 1mM pyruvate, and 20mM HEPES). Coverslips were then transferred to the lattice light-sheet microscope (LLSM) sample holder and inserted into the imaging chamber containing 6 ml of the later medium and 0.1 mg/ml of PKMR.

Previously described Hela cells expressing Rab5-eGFP CRISP genome edited SUM 159 breast carcinoma cell line, using eGFP-tagged *σ* unit of the AP2 complex (Adaptor Protein complex) were kindly provided by Dr. Tomas Kirchhausen (HMS, Boston, MS, USA) and CD-M6PR-EGFP expressing Hela cell line, by Dr Bernard Hoflack (TU-Dresden, Dresden, Germany). Human retinal pigment epithelial cells (RPE1) expressing mCherry-LifeAct were kindly provided by Dr Manuel Thery (Vignaud *et al*. 2012)^3^. All these cells were cultured, prepared and imaged with the LLSM as described above.

For STED imaging RPE1 cells were plated on glass bottom Petri dishes (*μ*-dish 35mm-High,1.5H, Ibidi, GmbH, Grafelfing, Germany). Microtubule labeling was performed as for LLSM imaging, while mitochondria were labeled by diluting 16000 × a 1mM stock solution of PKMITO^*TM*^ Orange [gifted by Dr. Z. Chen from Peking University and Genvivo Biotech, Beijing, PR of China] in DMEM/F12 medium for a final 62.5 nM, for 15 minutes at 37°C before image acquisition.

Previously acquired MFM datasets of U2OS10 cells transfected with TOM20 (translocase of outer mitochondrial membrane) fused to GFP (GFP-TOM20) (Hajj *et al*. 2014)^4^ were used in this study.

Previously acquired datasets of M10 cells, stably expressing Langerin tagged with eYFP, or Rab11a tagged with mCherry (Gidon *et al*. 2012)^5^) or transiently transfected with PhLuorin Transferrin-Receptor (TfR) (Boulanger *et al*. 2014)^6^) were also exploited in this study.

##### Microscopy

###### Lattice light-sheet microscopy (LLSM)

Lattice light-sheet microscopy was done on a commercialized version of a previously described setup Chen *et al*. 2012)^1^ from 3i (Denver, USA). Cells were scanned incrementally through a 20 *μ*m long light sheet in 600 nm steps using a fast piezoelectric flexure stage equivalent to ≃ 325 nm with respect to the detection objective and were imaged using a sCMOS camera (Orca-Flash 4.0; Hamamatsu, Bridgewater, NJ). Excitation was achieved with 560- or 642-nm diode lasers (MPB Communications) at 10% and 15% acousto-optic tunable filter transmittance with 50 mW and 100 mW respectively (initial box power) through an excitation objective (Special Optics 28.6 × 0.7 NA 3.74-mm water-dipping lens) and detected via a Nikon CFI Apo LWD 25 × 1.1 NA water-dipping objective with a 2.5 × tube lens with a final pixel size of 104 nm. Lattice light-sheet imaging was performed using an excitation pattern of outer NA equal to 0.55 and inner NA equal to 0.493. Composite volumetric datasets were obtained using ≃ 10 ms exposure/slice/channel at a time resolution of 2.2 s per total cell volume (about 60 planes). Fifty-six time points were acquired within 3 to 4 minutes (total raw data, only 1:45 min were analyzed for the movie). Acquired data were deskewed, a necessary step to realigned image frames, using LLSpy, a python library to facilitate lattice light sheet data processing (copyright to T. Lambert, Harvard Medical School, Boston, USA; https://github.com/tlambert03/LLSpy). Deskewed images are then considered as Raw images. Napari, a multi-dimensional image viewer for Python (https://github.com/napari/napari), was used for 3D rendering. Maximum intensity projections were generated using ImageJ/Fiji 1.53c (Schindelin *et al*., 2012)^7^. Intensity profiles were plotted using Matlab2019b. Related supplementary movies S1 and S2 were generated using Napari and napari-animation plugin (https://github.com/napari/napari-animation).

###### Multifocus microscopy (MFM)

MultiFocus microscopy (MFM) ^31^ allows simultaneous acquisition of a 3D stack of nine equally spaced focal 2D images with a frame rate up to 100 volumes per second, thus covering the whole volume of a nucleus or cell components in a single exposure ^31^. The set-up is based on a diffraction grating that forms multiple focus-shifted images followed by a chromatic correction module. This imaging technique is especially designed for high-speed imaging, and is able to capture the signal of weak fluorescent samples such as single fluorophores.

###### Stimulated emission depletion microscopy (STED)

Image acquisition was performed with a STEDYCON module (Abberior Instruments, Göttingen, Germany) mounted at the camera port TCS SP8 STED microscope (Leica, Manheim, Germany) with a HC PL APO C2S × 100 oil objective (1.4 NA) used in 2D mode. Depletion was performed with a 775 nm pulsed laser 1-7 ns at 80% (413 mW at the objective lens position). Labeled mitochondria and microtubules were imaged with excitation at a wavelength of 594 nm and 640 nm, respectively. Nominal laser powers were adjusted at 7% for the 594 nm laser (3.15 *μW* at the objective lens position) and at 10% for the 640 nm (26 *mW* at the objective lens position). The time-gated fluorescence detection was done on a detector single photon counting avalanche photodiode between 578-627 nm and 650-700 nm with a pinhole settled at 1.1 AU. Imaging was executed with the 9 lines accumulation. Pixel dwell time was 10 *μs* and a pixel size of 25 nm × 25 nm. Images were generated using ImageJ/Fiji 1.53c3. Line profiles plots were measured with ImageJ. Then line profiles were fitted to a Gaussian model using Matlab. Finally, Full width at half maximum (FWHM) is estimated from the fitting results.

###### Other Imaging modalities

Spinning disk confocal microscopy (CM) and Ring Total Internal Reflection microscopy (RING-TIRFM) were either used as described before (Gidon *et al*. 2012)^3^, (Pecot *et al*. 2018)^8^ or previously acquired datasets ((Boulanger *et al*. 2014)^4^; (Basset *et al*. 2015)^9^) reused, in this study.

#### D. Quantitative evaluation and method comparison on 2D artificially noisy and blurred fluorescence images

We demonstrate that SPITFIR(e) (based on Sparse Hessian Variation) provides the best PSNR (Peak Signal to Noise Ratio) values when the ground truth and the PSF are known, when compared to other competitive methods reported in Table 1. As the implementation in 3D of several regularizers in Table 1 is not always possible, we conducted fair experiments on 2D images shown in Supplementary Table 2. We included the performance of classical deconvolution methods such as Richardson-Lucy (RL), iterative constrained Tikhonov-Miller (ICTM), Gold-Meinel (GM)) ^32^ on the four tested images. The four ground-truth images in Supplementary Table 2 were normalized in the range [0, 1] and further blurred by considering a Gaussian point spread function (PSF) with different standard deviation values *σ*_*xy*_. A Gaussian noise with zero mean and variance *τ*^2^ was also added to these images in order to generate observed noisy and blurry data. Most of competing methods are known to perform better if the images are corrupted by Gaussian noise. Note that, in the case of RL algorithm, which is originally designed to deal with Poisson noise, the degraded images are re-scaled to the original dynamics of the underlying reference images. The RL deconvolution results are then re-normalized for a fair comparison with those obtained by the other methods. Moreover, before applying GM, the noisy images are smoothed with a Gaussian filter as pre-processing step because this algorithm assumes that the noise is negligible. In these experiments, we consider three different PSF sizes which correspond to *σ*_*xy*_ ∈ {1, 1.25, 1.5} and three distinct noise levels *τ* ∈ {0.01, 0.02, 0.04} (intensities are in the range [0, 1]).

The parameters involved in each method have been adjusted in order to get the highest PSNR values for each individual image in Supplementary Table 2. We can notice that the SHV-based deconvolution approach outperforms the competing methods in most cases (highest PSNR scores in 21 over 36 cases). In these cases, SV and TV-*L*_1_, which are respectively the second (12/36) and the third (7/36) position on the benchmark, achieve slightly lower PSNR scores while comparing with SHV. In contrast to these results, the non-regularization methods such as RL and GM algorithm are often ranked at the end of the benchmark. Last but not least, the GR measure that we thought a good alternative for TV, provides disappointing deconvolution outcomes in terms of PSNR values on artificially corrupted images.

In terms of visual assessment, SPITFIR(e) avoid over-smoothing and over-sharpening effect while preserving details and removing noise in the background, as illustrated on Image #1 in Supplementary Fig. 6. In Supplementary Fig. 6(**c**)-(**d**), we notice noise amplification with RL (**c**) and GM (**d**), resulting in the apparition of undesired high-intensity pixels (i.e. the “night sky” effect) and unrealistic reconstructed structures since the high-frequency components were not correctly restored. Unlike RL and GM algorithms, ICTM (**e**) and GraphNet (**h**) produce deconvolution results without noise amplification artifacts, but they tend to over-smooth (blur) image details due to the quadratic nature of Tikhonov-Miller penalty. In contrast to these over-smoothed results, those obtained with TV (**f**), TV-*L*_1_ (**g**) and SV (**l**) regularizers generate artificial sharper edges (staircasing effects), respectively. As depicted in Supplementary Fig. 6(**m**), HV which promotes piecewise linear reconstructions, provides visually more pleasant deconvolution results. Similar visual results are obtained with SPITFIR(e) (SHV, (**n**)), which enforce the co-localization of high-intensity pixels and non-zero directional derivatives, and allow to obtain non-zero regions with eventually high-magnitude gradients. It results in better reconstruction of the image foreground, in which large variation between pixels at the object center and those at the object boundary may occur. Regarding smooth-approximation based regularizations, TVH (**i**) and LSHV (**j**) provide results with restored details which are slightly sharper than those obtained with ICTM (**e**), but noise is not sufficiently removed. It not only disturbs the visual effect but also makes the detection of small or low-contrast objects more difficult. In our opinion, this issue is due to the smooth nature of the approximation for small values of directional derivatives, resulting in noisy reconstructed gradients. As consequence, the PSNR values obtained with these two regularizers are sharply inferior to those obtained with SV, HV and SHV. In the comparison with TVH (**i**) and LSHV (**j**), the smooth version of GR (**k**) yields visual similar deconvolution results. However, GR (**k**) suffer from severe “white pixel” artifacts in the background region. These “white pixels” are discarded to fairly compute the PSNR values. Overall, SHV (SPITFIR(e)) (**n**) provided the most satisfying results in this comparison experiment.

## Footnotes

1 B.-C. Chen *et al*., *Science*, 346(6208): 1257998 (2014). (doi:10.1126/science.1257998)

2 Z. Yang *et al*., *Chem. Sci*., 11:8506 (2020). (doi:10.1039/d0sc02837a)

3 Vignaud *et al*. J. Cell Sci. 125(9): 2134-2140 (2012). (doi: 10.1242/jcs.104901)

4 B. Hajj *et al*. Proc. Natl. Acad. Sci. USA, 111:17480-17485 (2014). (doi: 10.1073/pnas.1412396111)

5 A. Gidon *et al*., Traffic, 13(6):815-833 (2012). (doi:10.1111/j.1600-0854.2012.01354.x)

6 J. Boulanger *et al*., Proc. Natl. Acad. Sci. US, 111(8): 7164-17169 (2014). (doi:10.1073/pnas.1414106111)

7 Schindelin *et al*., *Nat. Methods*, 9(7), 676–682 (2012). (doi:10.1038/nmeth.2019)

8 T. Pécot *et al*., eLife, 7:e32311 (2018). (doi:10.7554/eLife.32311)

9 A. Basset *et al*., *IEEE Trans. Image Process*., 24(11):4512-4527 (2015). (doi:10.1109/TIP.2015.2450996)

## References

[1] Carlton, P. et al. Fast live simultaneous multi-wavelength 4-dimensional optical microscopy. Proc. Natl. Acad. Sci. USA 107, 16016–16022 (2010).

[2] Boulanger, J. et al. Patch-based nonlocal functional for denoising fluorescence microscopy image sequences. IEEE Trans. Medical Imaging 29, 442–454 (2010).

[3] Richardson, W. H. Bayesian-based iterative method of image restoration. J. Optical Society of America 62, 55–59 (1972). URL http://www.osapublishing.org/abstract.cfm?URI=josa-62-1-55.

[4] Lucy, L. B. An iterative technique for the rectification of observed distributions. The Astronomical J. 79, 745–754 (1974).

[5] Shepp, L. A. & Vardi, Y. Maximum likelihood reconstruction for emission tomography. IEEE Trans. Medical Imaging 1, 113–122 (1982).

[6] Sibarita, J.-B. Deconvolution Microscopy, 201–243 (Springer Berlin Heidelberg, Berlin, Heidelberg, 2005). URL https://doi.org/10.1007/b102215.

[7] van der Voort, H. T. M. & Strasters, K. C. Restoration of confocal images for quantitative image analysis. J. Microscopy 178, 165–181 (1995). URL http://dx.doi.org/10.1111/j.1365-2818.1995.tb03593.x.

[8] van Kempen, G. M. P., van der Voort, H. T. M., Bauman, J. G. J. & Strasters, K. C. Comparing maximum likelihood estimation and constrained Tikhonov-Miller restoration. IEEE Engineering in Medicine and Biology Magazine 15, 76–83 (1996).

[9] van Kempen, G. M. P., van Vliet, L. J., Verveer, P. J. & van der Voort, H. T. M. A quantitative comparison of image restoration methods for confocal microscopy. J. Microscopy 185, 354–365 (1997). URL http://dx.doi.org/10.1046/j.1365-2818.1997.d01-629.x.

[10] Dey, N. et al. ichardson-lucy algorithm with total variation regularization for 3d confocal microscope deconvolution. Microsc. Res. Tech. 69, 260–266 (2006).

[11] Hom, E. et al. Aida: An adaptive image deconvolution algorithm with application to multi-frame and three-dimensional data. J. Opt. Soc. Am. A 24, 1580–1600 (2007).

[12] Lefkimmiatis, S., Ward, J. P. & Unser, M. Hessian Schatten-norm regularization for linear inverse problems. IEEE Trans. Image Processing 22, 1873–1888 (2013).

[13] Ikoma, H., Broxton, M., Kudo, T. & Wetzstein, G. A convex 3d deconvolution algorithm for low photon count fluorescence imaging. Scientific Reports 8, 111489 (2018).

[14] Arigovindan, M. et al. High-resolution restoration of 3D structures from widefield images with extreme low signal-to-noise-ratio. Proc. Natl. Acad. Sci. USA 110, 17344–17349 (2013). URL http://www.pnas.org/content/110/43/17344.abstract. http://www.pnas.org/content/110/43/17344.full.pdf.

[15] Ng, B., Vahdat, A., Hamarneh, G. & Abugharbieh, R. Generalized sparse classifiers for decoding cognitive states in fMRI. In Wang, F., Yan, P., Suzuki, K. & Shen, D. (eds.) ProceedingsInt. MICCAI Workshop on Machine Learning in Medical Imaging (MLMI), 108–115 (Beijing, China, 2010).

[16] Kandel, B. M., Wolk, D. A., Gee, J. C. & Avants, B. Predicting cognitive data from medical images using sparse linear regression. In Gee, J. C., Joshi, S., Pohl, K. M., Wells, W. M. & Zöllei, L. (eds.) Int. Conf. Information Processing in Medical Imaging (IPMI 2013), 86–97 (Asilomar, CA, USA, 2013).

[17] Grosenick, L., Klingenberg, B., Katovich, K., Knutson, B. & Taylor, J. E. Interpretable whole-brain prediction analysis with GraphNet. NeuroImage 72, 304–321 (2013). URL http://www.sciencedirect.com/science/article/pii/S1053811912012487.

[18] Eickenberg, M., Dohmatob, E., Thirion, B. & Varoquaux, G. Grouping total variation and sparsity: Statistical learning with segmenting penalties. In Medical Image Computing and Computer-Assisted Intervention (MICCAI), 685–693 (Munich, Germany, 2015).

[19] Chan, T. F. & Esedoglu, S. Aspects of total variation regularized L1 function approximation. SIAM J. Applied Mathematics 65, 1817–1837 (2005). URL https://doi.org/10.1137/040604297.

[20] Michel, V., Gramfort, A., Varoquaux, G., Eger, E. & Thirion, B. Total variation regularization for fMRI-based prediction of behavior. IEEE Trans. Medical Imaging 30, 1328–1340 (2011).

[21] Zhao, W. et al. Sparse deconvolution improves the resolution of live-cell super-resolution fluorescence microscopy. Nat. Biotechnol. (2021).

[22] Afonso, M., Bioucas-Dias, J. & Figueiredo, M. An augmented lagrangian approach to the constrained optimization formulation of imaging inverse problems. IEEE Trans. Image Processing 20, 681–695 (2011).

[23] Boyd, S., Parikh, N., Chu, E., Peleato, B. & Eckstein, J. Distributed optimization and statistical learning via the alternating direction method of multipliers. Found. Trends Mach. Learn. 3, 1–122 (2007).

[24] Beck, A. & Teboulle, M. Fast iterative shrinkage-thresholding algorithm for linear inverse problems. SIAM J. Imaging Sciences 2, 183–202 (2009).

[25] Condat, L. A primal–dual splitting method for convex optimization involving lipschitzian, proximable and linear composite terms. J. Optimization Theory and Applications 158, 460–479 (2013). URL http://dx.doi.org/10.1007/s10957-012-0245-9.

[26] Condat, L. A generic proximal algorithm for convex optimization – application to total variation minimization. IEEE Signal Processing Letters 21, 985–989 (2014).

[27] Chen, B.-C. & et al. Lattice light-sheet microscopy: Imaging molecules to embryos at high spatiotemporal resolution. Nature 346, 1257998–1, 1257998–12 (2014).

[28] Yang, Z. et al. Cyclooctatetraene-conjugated cyanine mitochondrial probes minimize phototoxicity in fluorescence and nanoscopic imaging. Chem. Sci. 11, 8506–8516 (2020). URL http://dx.doi.org/10.1039/D0SC02837A.

[29] Renard, H. et al. Endophilin-a3 and galectin-8 control the clathrin-independent endocytosis of cd166. Nat. Commun 11 (2020).

[30] Lea, P. & Hollenberg, M. Mitochondrial structure revealed by high-resolution scanning electron microscopy. Am J Anat. 184 (1989).

[31] Hajj, B. et al. Whole-cell, multicolor superresolution imaging using volumetric multifocus microscopy. Proc. Natl. Acad. Sci. USA 111, 17480–17485 (2014).

[32] Sage, D. et al. DeconvolutionLab2: an open-source software for deconvolution microscopy. Methods 115, 28–41 (2017).

[33] Bruce, M. & Butte, M. Real-time gpu-based 3d deconvolution. Opt. Express 21, 4766–4773 (2013).

[34] Weigert, M., Schmidt, U. & et al., T. B. Content-aware image restoration: pushing the limits of fluorescence microscopy. Nat. Methods 15, 1090–1097 (2018).

[35] Kirshner, H., Aguet, F., Sage, D. & Unser, M. 3-D PSF fitting for fluorescence microscopy: Implementation and localization application. Journal of Microscopy 249, 13–25 (2013).

[36] Huang, X. et al. Fast, long-term, super-resolution imaging with hessian structured illumination microscopy. Nat. Biotechnol. 36, 451–459 (2018).

[37] Till, S., Roesch, A., Riedel, D. & Jakobs, S. Live-cell sted nanoscopy of mitochondrial cristae. Sci. Rep. 9 (2019).

[38] Yang, X. et al. Mitochondrial dynamics quantitatively revealed by sted nanoscopy with an enhanced squaraine variant probe. Nat. Commun 11 (2020).

[39] Mugnier, L., Fusco, T. & Conan, J.-M. Mistral: a myopic edge-preserving image restoration method, with application to astronomical adaptiveoptics-corrected long-exposure images. J. Opt. Soc. Am. A 21, 1841–1854 (2004).

[40] Soulez, F., Denis, L., Tourneur, Y. & Thiébaut, É. Blind deconvolution of 3D data in wide field fluorescence microscopy. In Int. Symp. Biomedical Imaging (ISBI) (Barcelone, Spain, 2012).

[41] Rudin, L. I., Osher, S. & Fatemi, E. Nonlinear total variation based noise removal algorithms. Physica D: Nonlinear Phenomena 60, 259–268 (1992). URL http://www.sciencedirect.com/science/article/pii/016727899290242F.

[42] Heideman, M., Johnson, D. & Burrus, C. Gauss and the history of the fast Fourier transform. IEEE ASSP Magazine 1, 14–21 (1984).

[43] Van Loan, C. Computational Frameworks for the Fast Fourier Transform (Society for Industrial and Applied Mathematics, 1992). URL http://epubs.siam.org/doi/abs/10.1137/1.9781611970999. http://epubs.siam.org/doi/pdf/10.1137/1.9781611970999.

[44] Frigo, M. & Johnson, S. G. FFTW: an adaptive software architecture for the FFT. In IEEE Int. Conf. Acoustics, Speech and Signal Processing (ICASSP), vol. 3, 1381–1384 (1998).

[45] Frigo, M. & Johnson, S. G. The design and implementation of FFTW3. Proceedings of the IEEE 93, 216–231 (2005).

[46] Beck, A. & Teboulle, M. Fast gradient-based algorithm for constrained total variation image denoising and deblurring problems. IEEE Trans. Image Processing 18, 2419–2434 (2009).

[47] Figueiredo, M., Bioucas-Dis, J. & Nowack, R. Majorization-minimization algorithms for wavelet-based image restoration. IEEE Trans. Image Processing 16, 2980–2991 (2007).

[48] Mercier, B. Lectures on Topics in Finite Element Solution of Elliptic Problems (Springer-Verlag Berlin Heidelberg, 1979).

[49] Eckstein, J. & Bertsekas, D. P. On the Douglas–Rachford splitting method and the proximal point algorithm for maximal monotone operators. Mathematical Programming 55, 293–318 (1992). URL http://dx.doi.org/10.1007/BF01581204.

[50] Combettes, P. L. Solving monotone inclusions via compositions of nonexpansive averaged operators. Optimization 53, 475–504 (2004).

[51] Combettes, P. L. & Wajs, V. R. Signal recovery by proximal forward-backward splitting. SIAM J. Multiscale Modeling and Simulation 4, 1168–1200 (2005). URL https://doi.org/10.1137/050626090.

[52] Combettes, P. L. & Pesquet, J.-C. Proximal Splitting Methods in Signal Processing, 185–212 (Springer New York, New York, NY, 2011). URL http://dx.doi.org/10.1007/978-1-4419-9569-8_10.

[53] Chambolle, A. & Pock, T. A first-order primal-dual algorithm for convex problems with applications to imaging. J. Math. Imaging and Vision 40, 120–145 (2011). URL http://dx.doi.org/10.1007/s10851-010-0251-1.

[54] Gennert, M. & Yuille, A. Determining the optimal weights in multiple objective function optmization. In Proc. IEEE Int. Conf. Comp. Vision (ICCV) (1988).

[55] Boulanger, J. et al. Fast high-resolution 3d total internal reflection fluorescence microscopy by incidence angle scanning and azimuthal averaging. Proc. Natl. Acad. Sci. USA 111, 17164–17169 (2014).

[56] Gidon, A. et al. Rab11A/MyosinVb/Rab11-FIP2 complex frames two late recycling steps of langerin from erc to plasma membrane. Traffic 13 (2012).

[57] Meinel, E. S. Origins of linear and nonlinear recursive restoration algorithms. J. Optical Society of America A 3, 787–799 (1986). URL http://josaa.osa.org/abstract.cfm?URI=josaa-3-6-787.

[58] Nikolova, M. Weakly constrained minimization: Application to the estimation of images and signals involving constant regions. J. Math. Imaging and Vision 21, 155–175 (2004). URL https://doi.org/10.1023/B:JMIV.0000035180.40477.bd.

[59] Good, I. J. & Gaskins, R. A. Nonparametric roughness penalties for probability densities. Biometrika 58, 255–277 (1971).

[60] Foi, A., Trimeche, M., Katkovnik, V. & Egiazarian, K. Practical Poissonian-Gaussian noise modeling and fitting for single-image raw-data. IEEE Transactions on Image Processing 17, 1737–1754 (2008).

[61] Combettes, P. L., Dũng,. & Vũ, B. C. Proximity for sums of composite functions. J. Math. Analysis and Applications 380, 680–688 (2011). URL http://www.sciencedirect.com/science/article/pii/S0022247X11002137.

[62] Combettes, P. L., Condat, L., Pesquet, J. C. & Vũ, B. C. A forwardbackward view of some primal-dual optimization methods in image recovery. In IEEE Int. Conf. Image Processing (ICIP), 4141–4145 (2014).

[63] Moreau, J. J. Proximité et dualité dans un espace hilbertien. Bulletin de la Société Mathématique de France 93, 273–299 (1965). URL http://www.numdam.org/item?id=BSMF_1965__93__273_0.

